# VPS9D1-AS1 overexpression amplifies intratumoral TGF-β signaling and promotes tumor cell escape from CD8^+^ T cell killing in colorectal cancer

**DOI:** 10.1101/2022.05.12.491618

**Authors:** Lei Yang, Xichen Dong, Zheng Liu, Jinjing Tan, Xiaoxi Huang, Tao Wen, Hao Qu, Zhenjun Wang

## Abstract

Efficacy of immunotherapy is limited in patients with colorectal cancer (CRC) because high expression of tumor-derived transforming growth factor (TGF)-β pathway molecules and interferon (IFN)-stimulated genes (ISGs) promotes tumor immune evasion. Here, we identified a long noncoding RNA (lncRNA), VPS9D1-AS1, which was located in ribosomes and amplified TGF-β signaling and ISG expression. We show that high expression of VPS9D1-AS1 was negatively associated with T lymphocyte infiltration in two independent cohorts of CRC. VPS9D1-AS1 served as a scaffolding lncRNA by binding with ribosome protein S3 (RPS3) and a competing endogenous RNA (ceRNA) to sponge miR-22-5p/514a-3p to increase the translation of TGF-β, TGFBR1, and SMAD1/5/9. VPS9D1-AS1 knockout downregulated OAS1, an ISG gene, which further reduced IFNAR1 levels in tumor cells. Conversely, tumor cells overexpressing VPS9D1-AS1 were resistant to CD8^+^ T cell killing and lowered IFNAR1 expression in CD8^+^ T cells. In a conditional overexpression mouse model, VPS9D1-AS1 enhanced tumorigenesis and suppressed the infiltration of CD8^+^ T cells. Treating tumor-bearing mice with antisense oligonucleotide drugs targeting VPS9D1-AS1 significantly suppressed tumor growth. Our findings indicate that the tumor-derived VPS9D1-AS1/TGF-β/ISG signaling cascade promotes tumor growth and enhances immune evasion and may thus serve as a potential therapeutic target for CRC.

## Introduction

Colorectal carcinoma (CRC) is a major cause of cancer-related death worldwide and shows a high propensity for metastatic dissemination (Siegel et al., 2020). Microsatellite-stable (MSS) CRC is regarded as immunologically ‘cold’, meaning that it is scarcely infiltrated by T cells and possibly nonimmunogenic and therefore unlikely to benefit from immune therapies (Guan et al., 2021). Thus, immune checkpoint blockade (ICB) is more effective in microsatellite instability-high (MSI-H) CRC but not in MSS (Liao et al., 2019). The lack of a DNA mismatch repair mechanism in MSI patients results in a higher tumor mutation burden and thus high neoantigen exposure that favors ICB (Lu et al., 2021). However, important immune features, including the degree of T cell infiltration and the differentiation or activation state of T cells, remain to be elucidated (Benci et al., 2019).

Effective ICB relies on CD8^+^ T cell infiltration in the tumor microenvironment (TME). However, advanced-stage tumor cells secrete high levels of transforming growth factor (TGF)-β to reduce the activity of intratumoral cytotoxic T lymphocytes (CTLs), thereby inhibiting their antitumor effector functions (Katlinski et al., 2017). As a result, solid tumors evade anticancer immunity by establishing immune-privileged niches in the TME (Chongsathidkiet et al., 2018). Increased TGF-β in the TME limits the adaptive immune responses by inhibiting T effector cell functions and ushering exhausted T cells to apoptosis (Tauriello et al., 2018) (Liu et al., 2020). The receptors of interferon (IFN) are found to regulate TGF-β signaling pathway and are associated with CD8^+^ T cell immunity (Mariathasan et al., 2018).

IFN signaling is essential for communication between tumor cells and their neighboring cells (Sistigu et al., 2014). Endogenous IFNs contribute to antitumor immunity by stimulating specific CD8α lineage dendritic cells (DCs) to cross-present antigens to CTLs (Katlinski et al., 2017) and provide a “third signal” to stimulate the clonal expansion of CD8^+^ T cells (Gracias et al., 2013). In contrast, high levels of tumor-derived IFN stimulating genes (ISGs) are associated with immunological resistance. For example, IFN alpha receptor (IFNAR)-1 knockout in mouse cancer cells provoked pronounced immune responses after ionizing radiation and the cancer cells were more susceptible to CD8^+^ T cell-mediated killing (Chen et al., 2019). In tumor cells, PDL1 expression is promoted by IFN-γ secretion, which results in tumor cells escaping immune elimination (Cerezo et al., 2018). Thus, the IFN pathway plays contradictory roles in tumor cells and T lymphocytes.

VPS9D1-AS1 (also known as MYU), a long noncoding RNA (lncRNA) that has been proven to be overexpressed in multiple types of cancers (Kawasaki et al., 2016; Tan and Yang, 2018; Wang et al., 2020), was identified as a target of Wnt/c-Myc signaling and exhibited pro-oncogenic roles (Kawasaki et al., 2016). Here, we first report that VPS9D1-AS1 is an essential lncRNA that decreases CD8^+^ T cell infiltration by enhancing TGF-β and ISG expression in CRC. In addition, we propose that VPS9D1-AS1 might serve as a drug target to enhance the efficacy of ICB treatment against CRC.

## Results

### Increased VPS9D1-AS1 levels are positively associated with TGF-β signaling in CRC tissues

To study the clinical relevance of VPS9D1-AS1 expression, we used RNAscope to evaluate VPS9D1-AS1 levels in two independent cohorts. The OUTDO cohort enrolled 158 CRC subjects, and the BJCYH cohort enrolled 49 CRC patients. The levels of VPS9D1-AS1 were significantly higher in cancer tissues than in normal intestinal epithelial tissues (Figures 1A∼B and S1A). The survival and pathological characteristic analyses demonstrated that the levels of VPS9D1-AS1 were significantly associated with overall survival (OS), TNM stage and tumor lymph node metastasis (Figures 1C and S1B). We further confirmed the overexpression of VPS9D1-AS1 in cancer tissues using qRT-PCR assays (Figures 1D∼E). The RNAscope results were validated by qRT-PCR analysis (Figure 1F).

**Fig. 1.**
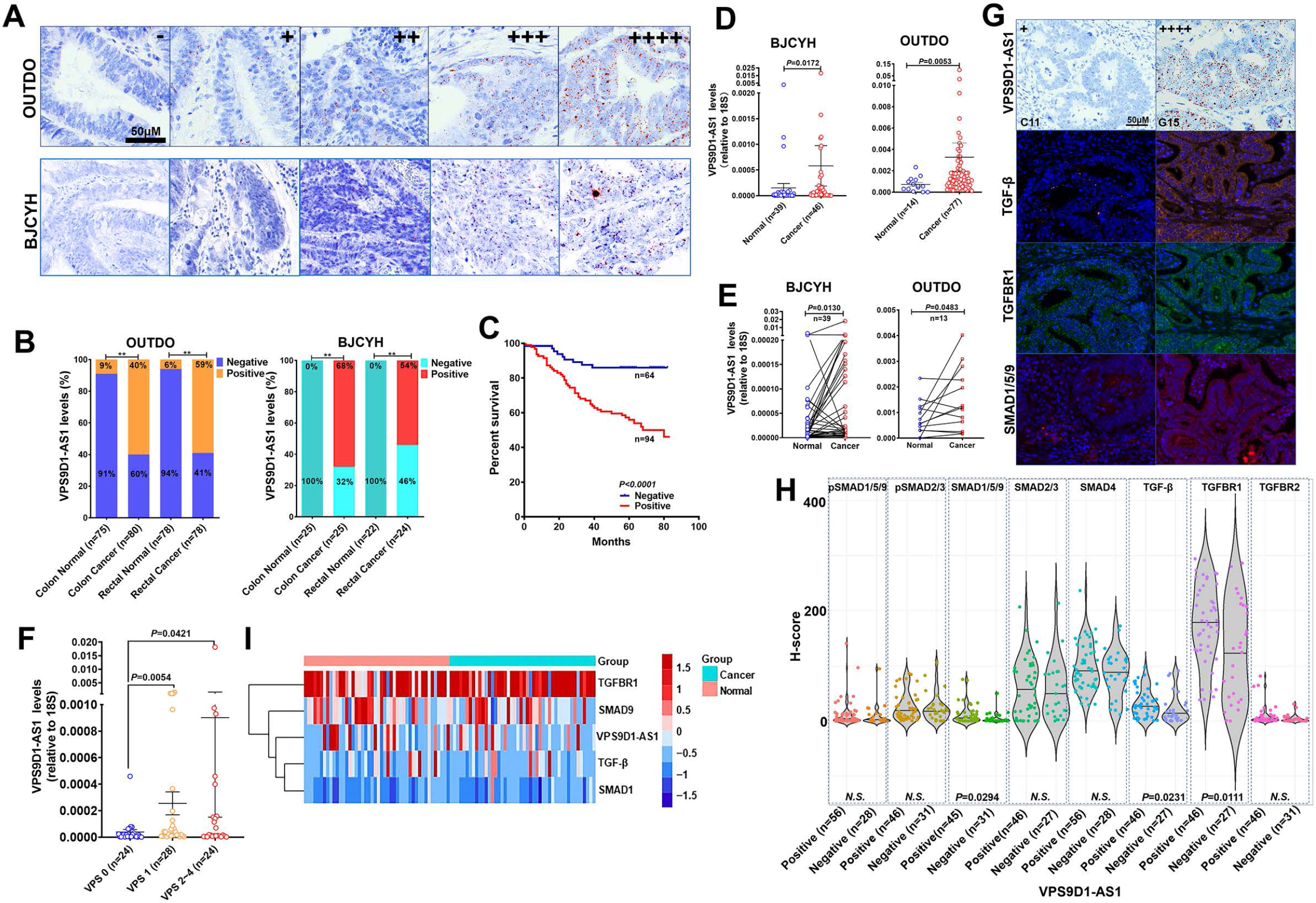
VPS9D1-AS1 is significantly upregulated in CRC and activates the TGF-β signaling pathway. (A) RNAscope stained VPS9D1-AS1 in CRC tissues that were enrolled in OUTDO and BJCYH cohorts. (B) Semiquantitative analyses of the levels of VPS9D1-AS1 in cancer and normal tissues of CRC patients. (C) Kaplan-Meier overall survival (OS) curves of VPS9D1-AS1-positive and VPS9D1-AS1-negative CRC patients. (D) qRT-PCR evaluation of the mRNA levels of VPS9D1-AS1. (E) Expression of VPS9D1-AS1 was compared in paired normal and cancer tissues. (F) The correlation between RNAscope and qRT-PCR assay results. (G) Representative pictures of VPS9D1-AS1-negative (+, C11) and VPS9D1-AS1-positive (G15, ++++) and multispectral fluorescence immunohistochemistry (mfIHC)-stained TGF-β, TGFBR1, and SMAD1/5/9 in the same CRC patients. (H) Integrative analysis of RNAscope and mfIHC data. (I) Heatmap clusters the mRNA levels of VPS9D1-AS1, TGF-β, TGFBR1, SMAD1, and SMAD9. *P* values were obtained by *chi*-square (B), log-rank test (C), unpaired *t* nonparametric test (D, F, H), and paired *t* test (E). Data are shown as data points with mean ± standard deviation of mean (SEM) (D), data points (E), data are depicted by violin and scatter plots with mean value (H). ** *P*<0.01. **Figure supplement 1**. Levels of VPS9D1-AS1 were not related to TGF-β signaling in cancer stromal cells.

Our previous study quantitatively investigated eight proteins (TGF-β, TGFBR1/2, SMAD1/5/9, pSMAD1/5/9, SMAD2/3, pSMAD2/3, and SMAD4) involved in TGF-β signaling by mfIHC staining (Yang et al., 2018; Yang et al., 2019). Because mfIHC and RNAscope assays were carried out on the same CRC tissue samples, we examined the relationships between VPS9D1-AS1 and TGF-β signaling. The protein levels of TGF-β signaling molecules were analyzed separately in the tumor and the cancer stroma. In tumor tissues, we found that higher levels of VPS9D1-AS1 were positively related to the expression of TGF-β, TGFBR1 and SMAD1/5/9 (Figures 1G∼H). We also detected the upregulation of mRNA encoding TGFBR1, SMAD1, and SMAD9 in tissue samples using qRT-PCR (Figures S1C∼D). However, there were no significant correlation at the mRNA level between VPS9D1-AS1 and the genes involved in the TGF-β signaling pathway (Figure 1I). In cancer stroma, VPS9D1-AS1 showed no effects on TGF-β signaling molecules (Figure S1E).

### Overexpression of VPS9D1-AS1 negatively associates with the levels of infiltrated cytotoxic T lymphocytes

To further explore the role of VPS9D1-AS1, we compared their levels in The Cancer Genome Atlas (TCGA) datasets that included 476 patients and their consensus molecular subtype (CMS) status (Guinney et al., 2015). Our analyses revealed that VPS9D1-AS1 was expressed predominantly in CMS2 patients (Figure S2A). The lymphocyte infiltration signature scores in CMS2 were significantly lower than those in CMS1, CMS3, and CMS4 (Figure S2B). Thus, we considered that the overexpression of VPS9D1-AS1 in CRC cells might be an important cause of the exclusion of T-infiltrating lymphocytes (TIL) from the TME.

To validate this hypothesis, we evaluated the levels of TILs in CRC tissue samples. In the OUTDO cohort, an mfIHC assay was carried out to calculate the percentages (%) of T lymphocytes in the total cancerous and cancer stromal cells, which represent the levels of T cell infiltration. TILs included CD4^+^, CD8^+^, and FOXP3^+^ T cells, and all subsets were significantly reduced in the cancerous tissues compared to the cancer stromal tissues (Figures 2A and S2C). In the BJCYH cohort, IHC assays demonstrated that the levels of CD8^+^ T cells were decreased while FOXP3^+^ T cells were increased in cancer tissues in comparing with matched normal tissues (Figure 2B and S2D).

**Fig. 2.**
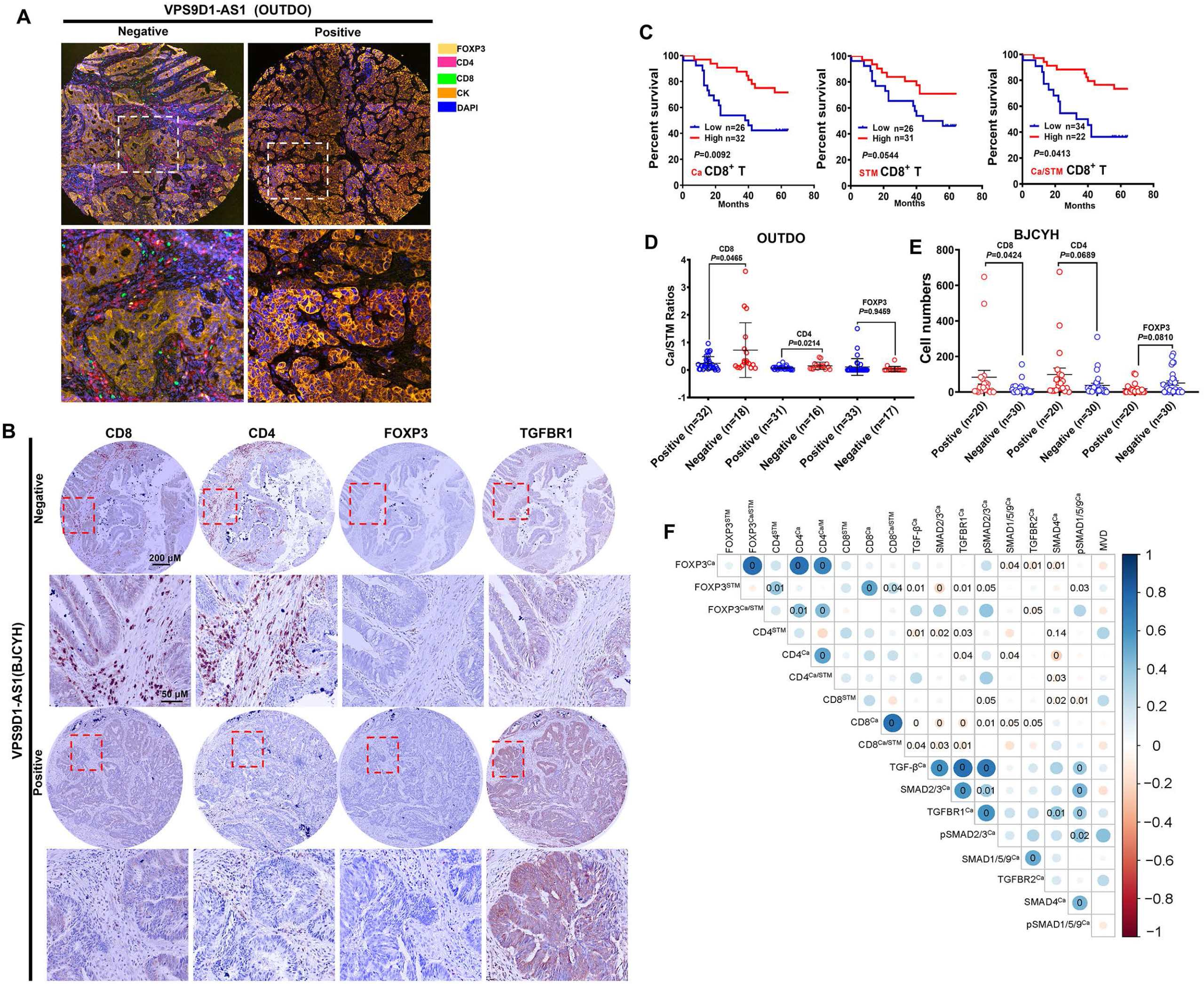
VPS9D1-AS1 is associated with reduced T lymphocyte infiltration. (A) Representative pictures of T cell infiltration (CD4, CD8, FOXP3) in CRC tissues for VPS9D1-AS1 quantification. Tumor cells are marked by cytokeratin (CK). (B) Representative pictures of CD8^+^, CD4^+^, FOXP3^+^ T cells and TGFBR1 stained by IHC in the BJCYH cohort. (C) The OS curves depicting the percentage of surviving CRC patients stratified by the levels of CD8^+^ T cell infiltration in cancerous (Ca) tissues, cancer stroma (STM), and the Ca/STM ratio. (D) Ca/STM ratios of CD4^+^, CD8^+^, and FOXP3^+^ T cells were calculated to identify the difference between VPS9D1-AS1 negative and positive populations in the OUTDO cohort. (E) The numbers of CD4^+^, CD8^+^, and FOXP3^+^ T cells in cancer tissues of BJCYH cohort were compared between VPS9D1-AS1 negative and positive tissues. (F) *Pearson* correlation analyses investigated the relationships between TILs and TGF-β signaling in cancer tissues. Eight proteins levels were investigated by mfIHC assays in same samples and fluorescence intensity of each protein level was transferred into quantitative data for *Pearson* correlation analyses. *P* values were obtained by log-rank test (C), unpaired *t* nonparametric test (D, E), and *Pearson* correlation test (F). Data are shown by mean ± SEM (D, E). **Figure supplement 2**. Integrative analysis of the relationship between VPS9D1-AS1, TGF-β signaling and TILs.

In the OUTDO cohort, the percentages of CD4^+^ T and FOXP3^+^ T cells in cancerous tissues (Ca) and cancer stroma (STM) and their ratios (Ca/STM) did not show any statistically significant relationship with overall survival (OS) (Figure S2E). In contrast, our analyses revealed that the levels of CD8^+^ T cells in Ca and STM and the Ca/STM ratio were significantly associated with OS (Figure 2C). We next tried to investigate the relationship between VPS9D1-AS1 and TILs and found that the levels of TILs were not significantly different between patients with low levels of VPS9D1-AS1 and those with high levels of VPS9D1-AS1 (Figure S2F). Interestingly, the levels of VPS9D1-AS1 were related to the Ca/STM ratios of CD4^+^ and CD8^+^ T cells (Figure 2D), suggesting that VPS9D1-AS1 prevented T cells from entering cancer tissues. In the BJCYH cohort, the levels of VPS9D1-AS1 were also negatively associated with TILs (Figure 2E). We further performed a *Pearson* correlation analysis to explore the relationships between the levels of TGF-β signaling molecules and TILs in both tumor and cancer stromal tissues. In tumor cells, the protein levels of TGF-β, TGFBR1, SMAD2/3 and pSMAD2/3 were negatively associated with the Ca/STM ratio of CD8^+^ T cells, and the levels of SMAD4 were negatively associated with the Ca/STM ratio of CD4^+^ T cells (Figure 2F), which is consistent with the role of TGF-β signaling in suppressing TILs. On the other hand, the protein levels of TGF-β, SMAD2/3, TGFBR1 and pSMAD1/5/9 in cancer stromal cells were positively associated with FOXP3^+^ T cell infiltration but negatively associated with CD8^+^ T cell infiltration (Figure S2G). Together, these results suggested that high expressions of VPS9D1-AS1 were positive associated with TGF-β signaling and negative associated with the levels of infiltrated CD8^+^ cytotoxic T cells.

### VPS9D1-AS1 is a tumor driver and positively regulates TGF-β signaling

First, we determined the levels of VPS9D1-AS1 in 16 cell lines and found that CRC cells expressed higher levels of VPS9D1-AS1 than other cells (Figure 3A). We designed four small guide RNAs targeting VPS9D1-AS1 (sgVPS) and used CRISPR/Cas9 to generate stable knockout (KO) CRC cell lines (Figures 3B and S3A∼B). VPS9D1-AS1 KO significantly downregulated TGF-β, TGFBR1, and SMAD1/5/9, did not affect SMAD2/3 and SMAD4, and increased SMAD6 expression, which acts as a negative regulator of TGF-β signaling (Figures 3C and S3C∼D). Furthermore, inferring RNA (siRNA) was used to disrupt the expression of VPS9D1-AS1 in HCT116 cells. We confirmed that VPS9D1-AS1 knockdown (KD) decreased TGF-β, TGFBR1, and SMAD1/5/9 expression (Figure S3E∼F). In contrast, ectopically overexpressed VPS9D1-AS1 increased the expression of TGF-β, TGFBR1, and SMAD1/5/9 and conferred resistance to the inhibition by SB431542 (a small molecule inhibitor of TGFBR1) (Figures S3G∼H). Moreover, VPS9D1-AS1 KD (both sgRNA and siRNA) had no impact on the mRNA expression of *TGF-β, TGFBR1*, and *SMAD1, ∼5, ∼9* (Figures S4A∼B).

**Fig. 3.**
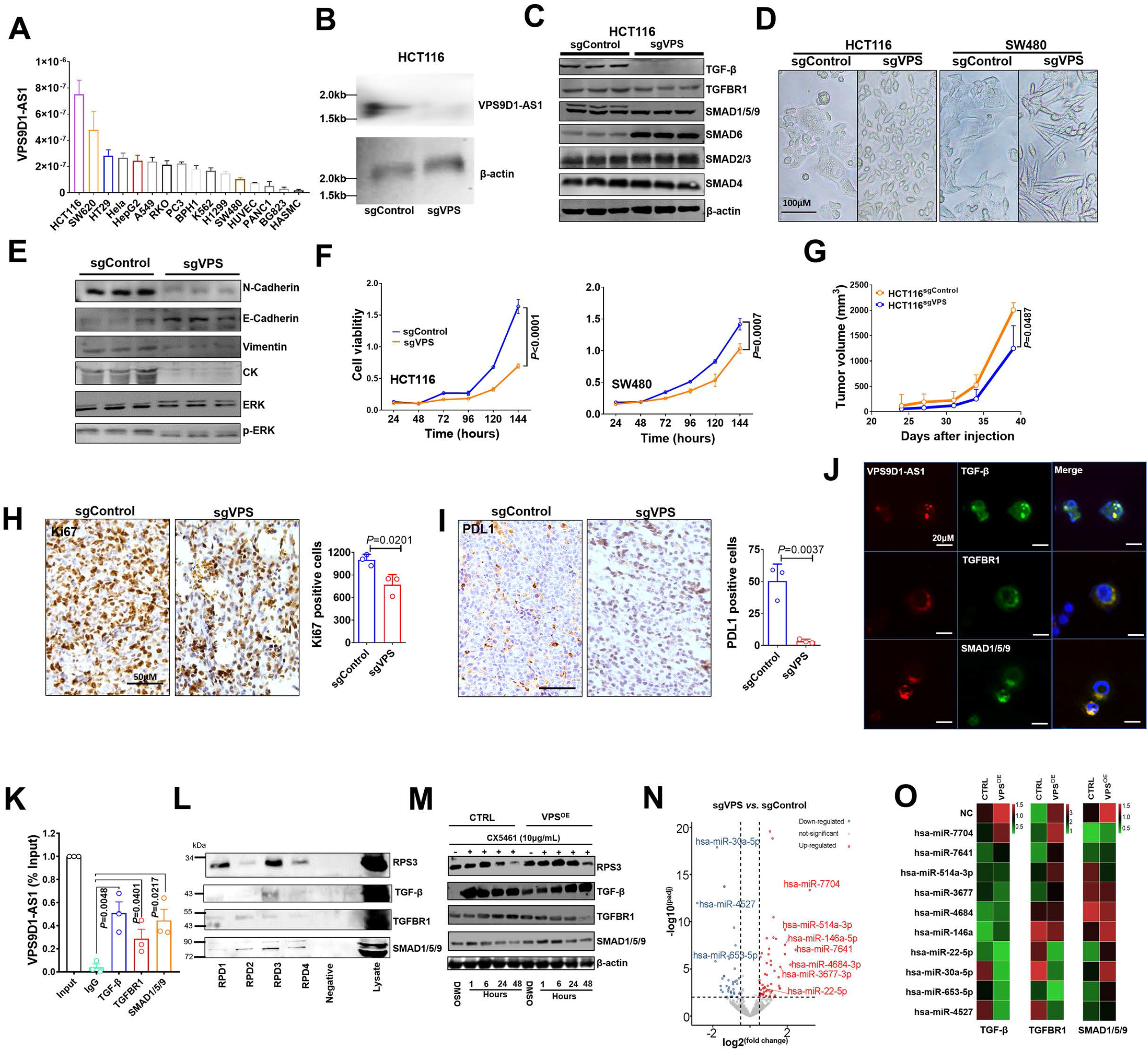
VPS9D1-AS1 controls TGF-β signaling and drives cell proliferation and metastasis. (A) The levels of VPS9D1-AS1 were determined by qRT-PCR in 16 cell lines. (B) Northern blotting validated the knockout (KO) of VPS9D1-AS1. (C) Western blotting measured the levels of proteins involved in TGF-β signaling. (D) Representative pictures show the cell morphologies of HCT116 and SW480 cell lines. (E) Western blotting quantified the levels of proteins involved in the EMT and ERK pathways (the same reference gene was used as in Figure 3C). (F) The proliferation of HCT116/SW480 sgControl and sgVPS cells were determined by CCK-8 assays. (G) Proliferation of xenograft tumors derived from HCT116 sgControl and sgVPS cells. (H) IHC determined the levels of Ki67 and (I) PDL1 in xenograft tissues. (J) RNA fluorescence *in situ* hybridization (FISH)-immunofluorescence (IF) and (K) RNA immunoprecipitation (RIP) assays showed the interaction between VPS9D1-AS1 and proteins that included TGF-β, TGFBR1, and SMAD1/5/9. (L) RNA pulldown-Western blotting assays detected the interaction between VPS9D1-AS1 and the intended proteins. (M) Western blotting determined the changes in RPS3, TGF-β, TGFBR1, and SMAD1/5/9 in HCT116 CTRL and VPS9D1-AS1 (VPS)-overexpressing (OE) cells treated with CX5461. RPD, RNA pull down probe. (N) Volcano plot displaying the differential expression of miRNAs between HCT116 sgControl and sgVPS cells. (O) Heatmaps showing the levels of TGF-β, TGFBR1, and SMAD1/5/9 in HCT116 CTRL and VPS OE cells transfected with mimics of 10 miRNAs. *P* values were obtained by two-way ANOVA (F, G) and paired or unpaired *t* tests (H, I, K). Data are shown as the mean ± SEM (F, G, H, I, K). **Source data1**. TGF-β, TGFBR1, SMAD4, SMAD1/5/9, SMAD6, SMAD2/3 and β-actin western-blot for Fig. 3C. **Source data 2**. N-Cadherin, E-Cadherin, Vimentin, CK, ERK and p-ERK western-blot for Fig. 3E. **Source data 3**. SMAD1/5/9, PRS3, TGF-β and TGFBR1 RNA-pull-down-western-blot for Fig. 3L. **Source data 4**. SMAD1/5/9, PRS3, TGF-β and TGFBR1 western-blot for Fig. 3M. **Figure supplement 3**. VPS9D1-AS1 activated TGF-β signaling. **Figure supplement 4**. VPS9D1-AS1 regulated TGF-β signaling and promoted tumor proliferation and migration. **Figure supplement 5**. VPS9D1-AS1 functions as the scaffolding lncRNA and ceRNA.

We next asked whether there was feedback between VPS9D1-AS1 and TGF-β signaling. Human recombinant (h)TGF-β protein and SB431542 were used to treat SW480 and HCT116 cells. These treatments had no significant effect on VPS9D1-AS1 levels in these cell lines (Figures S4C∼D). On the other hand, the downregulation of TGF-β, TGFBR1, and SMAD1/5/9 by siRNAs reduced the levels of VPS9D1-AS1 by 40%∼60% compared with the controls (siNC) (Figure S4E). These results indicated that loss of the endogenous TGF-β signaling molecules altered the expression of VPS9D1-AS1 through a feedback loop. However, manipulating the TGF-β signaling pathway with exogenous stimuli had no effects.

We next addressed the oncogenic roles of VPS9D1-AS1 such as promoting cell proliferation, migration, and clone formation. First, we found that stable VPS9D1-AS1 KO cells exhibited morphological changes (Figure 3D) and decreased clone formation capacity (Figure S4F). VPS9D1-AS1 KO significantly inhibited cell proliferation and migration (Figures 3F and S4G). Consistent with these observations, mechanistic analyses revealed that VPS9D1-AS1 KO reduced the levels of ERK, pERK, N-cadherin, vimentin, and cytokeratin but increased the level of E-cadherin (Figures 3E and S4H). In xenograft models, VPS9D1-AS1 KO significantly reduced the tumor volumes compared with the controls and significantly decreased the Ki67 and PDL1 levels in xenograft tumors (Figures 3G∼I and S4I). Specifically, SW480 VPS9D1-AS1 KO cells did not form xenograft tumors in mice (Figure S4I). Taken together, these findings support the notion that VPS9D1-AS1 acts as the driver of tumor progression by activating the ERK and EMT pathways.

### VPS9D1-AS1 scaffolds TGF-β signaling-related proteins and acts as a ceRNA to sponge miRNAs targeting TGF-β signaling

We predicted that VPS9D1-AS1 might act as scaffolding lncRNA in tumor cells. To validate this hypothesis, RNA FISH-immunofluorescence (IF) assays were conducted and the colocalization of VPS9D1-AS1, TGF-β, TGFBR1 and SMAD1/5/9 was confirmed in SW480 cells (Figure 3J). RNA immunoprecipitation (RIP) assays further showed that VPS9D1-AS1 directly bound to TGF-β, TGFBR1, and SMAD1/5/9 (Figure 3K). To map the protein binding regions in VPS9D1-AS1, we synthesized four biotinylated RNA probes targeting different regions of the VPS9D1-AS1 transcript (Figure S5A). These probes were used to perform RNA pulldown assays. As shown in Figure 3L, the interaction between VPS9D1-AS1 and TGF-β, TGFBR1 and SMAD1/5/9 proteins were validated.

We next predicted the subcellular localization of VPS9D1-AS1 by lncLocator (Lin et al., 2021) and found that most transcripts of VPS9D1-AS1 were localized in ribosomes (Figure S5B). Our RNA pulldown assay also proved that VPS9D1-AS1 bound with RPS3, one of the proteins constituting the small ribosomal subunit (Figure 3L). Thus, we sought to determine whether preventing ribosome biogenesis plays a role in regulating the translation of TGF-β, TGFBR1, SMAD1/5/9 (Devlin et al., 2016). We found that VPS9D1-AS1 overexpression (OE) prevented RPS3 degradation caused by CX5461 (an inhibitor of RNA polymerase I transcription of ribosomal RNA genes) treatment. However, CX5461 treatment immediately increased the levels of TGF-β, which declined over time. In VPS9D1-AS1 OE cells, the levels of TGFBR1 and SMAD1/5/9 were decreased after treatment with CX5461, but the TGF-β levels did not decrease following the degradation of RPS3 (Figure 3M). These findings suggest that VPS9D1-AS1 scaffolds the TGF-β protein and regulates its translation in ribosomes. We next identified 47 upregulated miRNAs and 40 downregulated miRNAs caused by VPS9D1-AS1 KO (Figure 3N). In HCT116 cells, the top 10 miRNAs regulated by VPS9D1-AS1 were selected, their mimics were transiently transfected, and the levels of TGF-β, TGFBR1, and SMAD1/5/9 were determined (Figures 3O and S5C). Among them, hsa-miR-22-5p, hsa-miR-514a-3p, and hsa-miR-146a were found to inhibit the expression of TGF-β, while hsa-miR-22-5p reduced the expression of TGFBR1 and SMAD1/5/9. Hsa-miR-7704 and hsa-miR-22-5p decreased the levels of SMAD1/5/9. We thus selected these four miRNAs to evaluate their roles in RKO and SW480 cells. VPS9D1-AS1 KO increased the expression of hsa-miR-22-5p, hsa-miR-514a-3p, and hsa-miR-7704. In contrast, VPS9D1-AS1 OE resulted in the downregulation of hsa-miR-22-5p and hsa-miR-514a-3p (Figure S5D). In agreement with these findings, we also observed that RPS3 interacted with hsa-miR-22-5p and hsa-miR-514a-3p (Figure S5E). Taken together, we concluded that VPS9D1-AS1 acted as a ceRNA to sponge hsa-miR-22-5p and hsa-miR-514a-3p, which targeted the mRNAs of *TGF-β, TGFBR1*, and *SMAD1/5/9*.

### IFN signaling activation induced by VPS9D1-AS1 expression acts downstream of TGF-β signaling

RNA sequencing was performed to identify the mRNAs differentially expressed between HCT116 sgControl and sgVPS cells. A total of 705 differentially expressed genes were identified, which included 203 upregulated genes and 502 downregulated genes (Figures 4A and S6A). VPS9D1-AS1 KO significantly inactivated IFNα/β signaling and the cell death pathway as well as immune system processes (Figures 4B and S6B∼C). Seventeen genes involved in IFNα/β signaling were validated in HCT116, RKO, and SW480 cells. IFI27 and OAS1 were the most significantly downregulated genes upon VPS9D1-AS1 KO (Figures 4C and S6D). Analyses in TCGA datasets demonstrated that OAS1 and IFI27 were significantly overexpressed in CRC cancer tissues (Figure S6E). We also analyzed the mRNA expression of OAS1 and IFI27 in tissue samples from 26 CRC cancer tissues and 10 normal colon tissues (Figure 4D). *Pearson* correlation analysis revealed that the levels of OAS1, but not IFI27, were significantly related to VPS9D1-AS1 levels (Figure 4E).

**Fig. 4.**
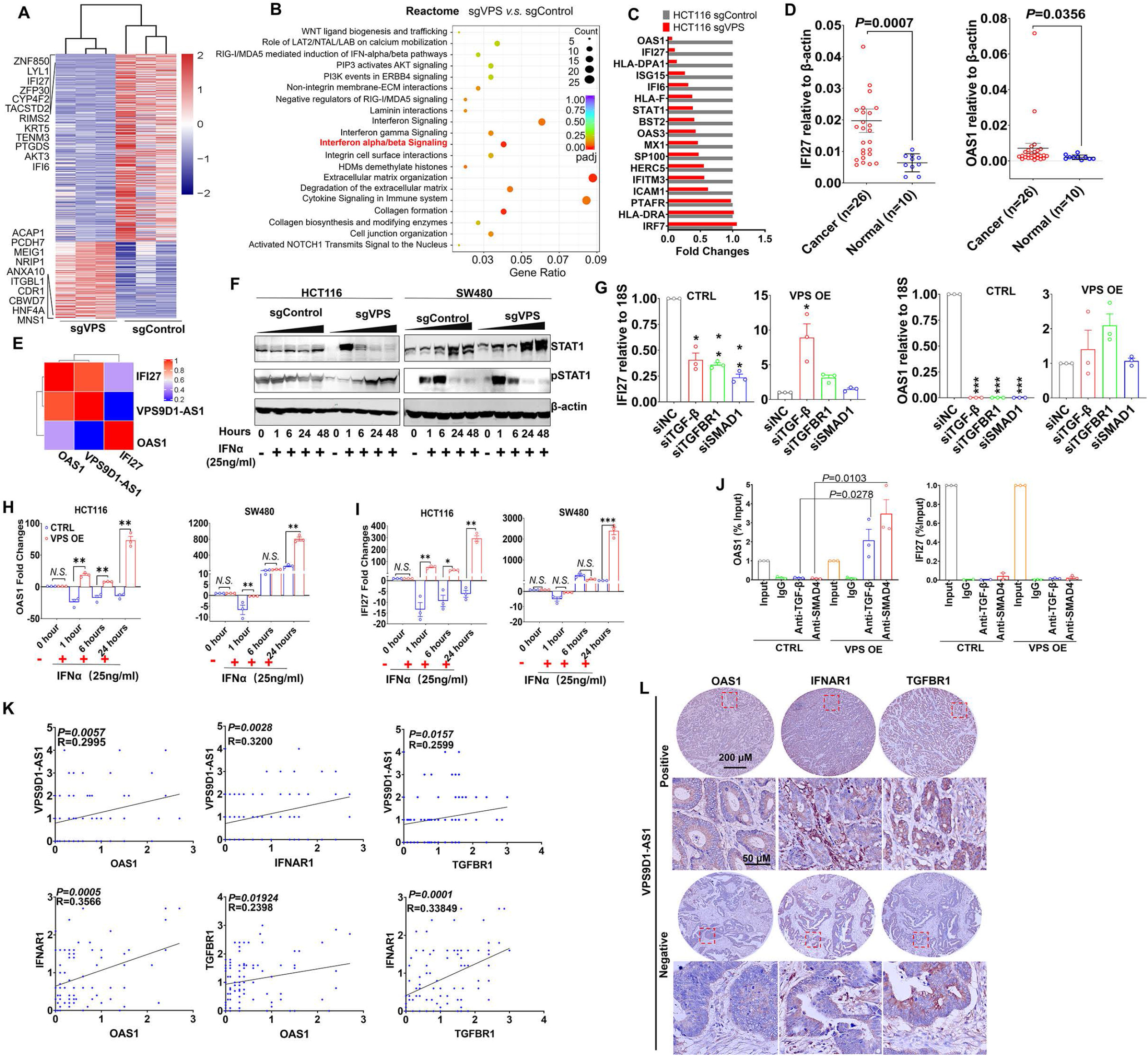
VPS9D1-AS1 regulates interferon signaling. (A) Heatmap illustrating the results of RNA sequencing of the genes regulated by VPS9D1-AS1. (B) VPS9D1-AS1 regulated the pathways associated with interferon signaling. (C) Differential expression of 17 genes in the IFNα/β signaling pathway was validated in HCT116 cells. (D) mRNA levels of IFI27 and OAS1 were determined by qRT-PCR, and (E) their relationships with VPS9D1-AS1 were calculated with *Pearson* correlation analysis. (F) Effect of VPS9D1-AS1 on STAT1 pathway activation induced by human recombinant IFNα. (G) VPS9D1-AS1 OE increased the expression of IFI27 and OAS1 through activated TGF-β signaling. (H) OAS1 and (I) IFI27 levels exhibited disparate changes upon IFNα stimulation in VPS9D1-AS1 OE cells and CTRL cells. (J) Chromatin immunoprecipitation (ChIP) assays demonstrated the interactions between TGF-β and SMAD4 and the promoter regions of OAS1 and IFI27. (K) *Pearson* correlation analyses investigated the relationships among VPS9D1-AS1, OAS1, IFNAR1, and TGFBR1 in CRC tissues. (L) IHC assays showed the levels of OAS1, IFNAR1, and TGFBR1 in patients with negative or positive expression of VPS9D1-AS1. *P* values were obtained by unpaired *t* test (D, J), two-way ANOVA (G, H, I) and *Pearson* correlation (K). Data are shown as the mean ± SEM (D, G, H, I, J). \**P*<0.05, ** *P*<0.01, *** *P*<0.001. *N*.*S*., not significant. **Source data 1**. SMAD1/5/9, PRS3, TGF-β and TGFBR1 western-blot for Fig. 4F **Figure supplement 6**. VPS9D1-AS1 plays a role on interferon signaling.

STAT1 is a well-known transcription factor activated by various ligands, including IFNα (Cerezo et al., 2018). After hIFNα stimulation, we found that VPS9D1-AS1 KO resulted in the downregulation of STAT1 but enhanced the levels of pSTAT1 at 1 hour (Figure 4F). This effect disappeared at 6, 24, and 48 hours, indicating that VPS9D1-AS1 KO regulated STAT1 phosphorylation and induced the immediate effect. When the cells were treated with hTGF-β, the phosphorylation of STAT1 induced by hIFNα stimulation was inhibited (Figure S6F). We further performed siRNA-mediated knockdown of TGF-β, TGFBR1, and SMAD1 and confirmed that blocking TGF-β signaling reduced OAS1 and IFI27 expression. The expression levels of OAS1 (but not IFI27) were more significantly affected. In contrast, VPS9D1-AS1 OE restored OAS1 and IFI27 expression (Figure 4G). Surprisingly, OAS1 and IFI27 mRNA levels were significantly reduced in VPS9D1-AS1 OE cells, although VPS9D1-AS1 was stably upregulated by ∼180.40 times in HCT116 cells and ∼42.28 times in SW480 cells (Figures S6G∼H). However, when hIFNα stimulation significantly increased OAS1 and IFI27 in VPS9D1-AS1 OE cells (Figures 4H∼I). These results led us to hypothesize that OAS1 activated IFN signaling through a feedback loop that was dependent on VPS9D1-AS1.

SMAD4 and TGF-β are noncanonical and canonical SMAD pathway cytokines, respectively, and they enter the nucleus and act as transcription factors (Derynck et al., 2021). We confirmed that both TGF-β and SMAD4 antibodies immunoprecipitated the promoter regions of both the *OAS1* (−77∼+284) and *IFI27* (−1933∼-1843) genes (Figure S6I). Importantly, VPS9D1-AS1 OE significantly enhanced the binding between TGF-β/SMAD4 and the promoter regions of *OAS1* but failed to enhance this binding in the *IFI27* promoter region (Figure 4J). IHC analysis indicated that the levels of OAS1, IFNAR1, and TGFBR1 were consistently elevated in CRC tissue samples that had positive VPS9D1-AS1 expression (Figure 4K∼L).

### VPS9D1-AS1 mediates the crosstalk between tumors and T cells depending on IFNAR1 expression

The receptor for IFNα/β is one of the downstream targets of ISGs. When we examined the changes in IFNAR1 on the surface of HCT116 cells by flow cytometry (FCM), we found that VPS9D1-AS1 KO reduced the expression of IFNAR1 (Figure 5A). Conversely, VPS9D1-AS1 OE significantly upregulated the expression of IFNAR1 in tumor cells (Figure 5B). To further delineate the upstream pathway essential for IFNAR1 upregulation in tumor cells, we applied the CRISPR/Cas9 technique to abolish OAS1 expression (Figure S7A). Our results demonstrated that the deletion of OAS1 decreased the expression of IFNAR1, although VPS9D1-AS1 OE cells were resistant to this effect (Figure 5B). In addition, the deletion of OAS1 inhibited STAT1 and TGF-β expression (Figure S7A).

**Fig. 5.**
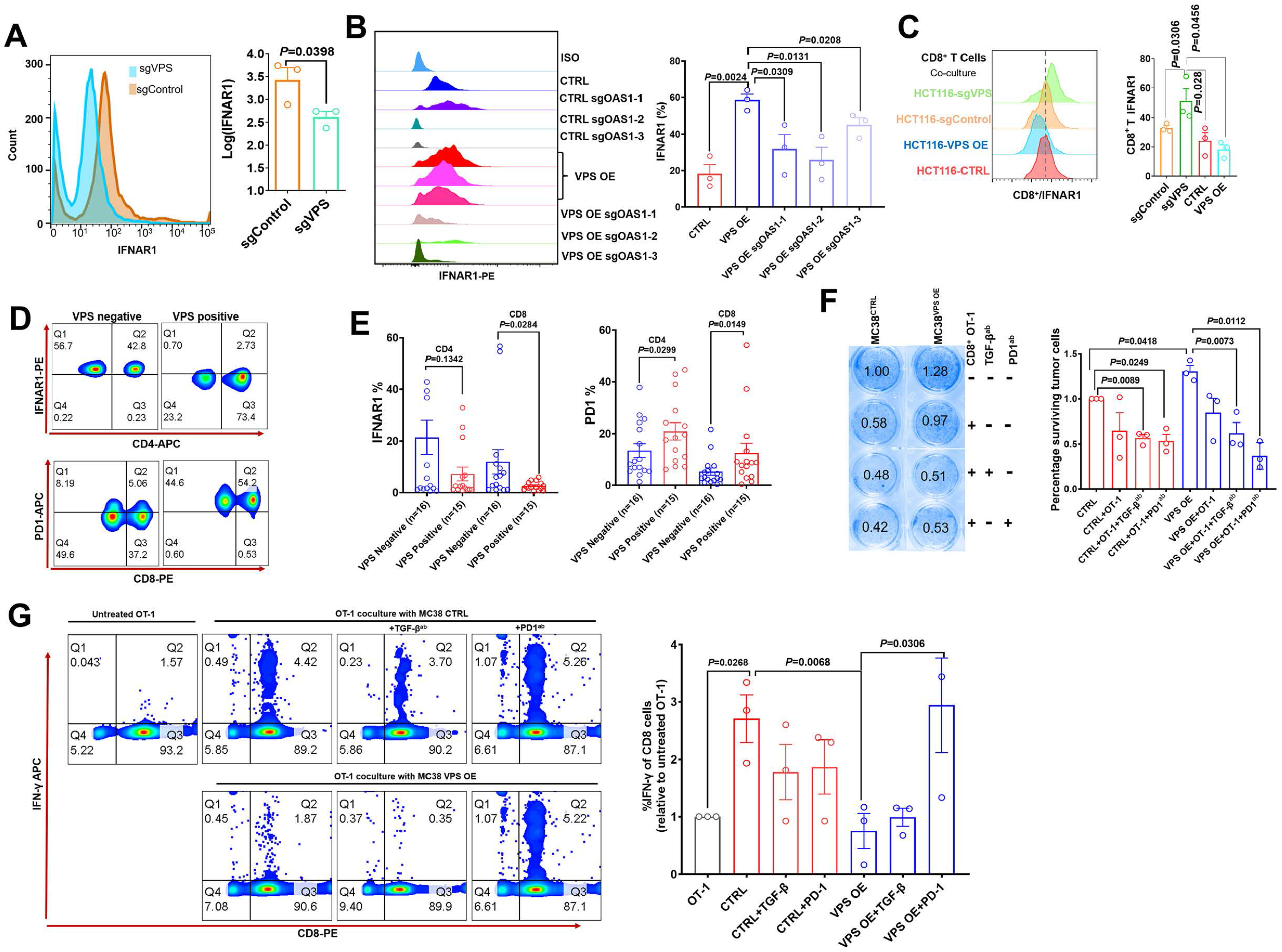
IFNAR1 mediates crosstalk between T cells and cancer cells. (A) Flow cytometry (FCM) revealed the decrease in IFNAR1 after VPS9D1-AS1 KO in HCT116 cells and (B) in HCT116 sgControl and sgVPS cells after CRISPR/Cas9-mediated inhibition of OAS1. Experiments were repeated three times. (C)Levels of IFNAR1 on the surface of CD8^+^ T cells cocultured with HCT116 CTRL, VPS OE, sgControl and sgVPS cells was detected by FCM in three independent assays. (D), (E) FCM determined the levels of IFNAR1 and PD1 in CD4^+^ and CD8^+^ T cells in peripheral blood. (F) T cell cytotoxicity assays against CTRL and VPS9D1-AS1-OE MC38-OVA cell lines by OT-1 CD8^+^ T cells. (G) FCM determined the IFN-γ levels in OT-1 CD8^+^ T cells after exposure to CTRL and VPS-OE MC38 cells. Antibodies against TGF-β and PD1 were added to the medium. The experiments were repeated three times. *P* values were obtained by unpaired *t* test (A, B, C, E, F, G). Data are shown as the mean ± SEM (A, B, C, E, F, G). **Figure supplement 7**. IFNAR1 mediates crosstalk between T cells and cancer cells.

To investigate the interaction of T cells and tumor cells, we prepared T cells from the peripheral blood of healthy donors. CD8^+^ T cells and total lymphocytes were separately cultured and primed with human interleukin 2 and antibodies against CD3 and CD28. In vitro-primed T cells were cultured with HCT116 sgControl and sgVPS cell lines. Coculture with VPS9D1-AS1 KO cells increased IFNAR1 expression in CD8^+^ and CD4^+^ T cells (Figures 5C and S7B). We also found that higher levels of VPS9D1-AS1 in cancer tissues were negatively associated with the expression levels of IFNAR1 but positively associated with the expression levels of PD1 in peripheral blood CD4/8^+^ T cells from CRC patients (Figures 5D∼E).

We further developed a T cell cytotoxicity assay using MC38-OVA cells. Our models successfully demonstrated that primed CD8^+^ T cells from OT-1 mice suppressed the proliferation of tumor cells (Figure S7C). On the other hand, VPS9D1-AS1 OE reduced the cytotoxicity of activated OT-1 CD8^+^ T cells (Figure 5F). Interestingly, both anti-TGF-β and anti-PD1 antibodies enhanced the cytotoxicity of CD8^+^ T cells in killing VPS9D1-AS1 OE cells (Figure 5F). FCM analysis showed that CD8^+^ T cells secreted more IFN-γ once they contacted tumor cells. However, VPS9D1-AS1 OE tumor cells inhibited CD8^+^ T cells from secreting IFN-γ. Furthermore, neutralizing antibodies against PD1 restored IFN-γ secretion by CD8^+^ T cells (Figure 5G).

These data support the idea that VPS9D1-AS1 upregulates OAS1 by enhancing TGF-β signaling derived from cancer cells to protect themselves from T cell-mediated cytotoxicity through regulation of IFNAR1.

### Upregulation of VPS9D1-AS1 inhibits antitumor immune cell infiltrations in immune-competent mice

The human *VPS9D1-AS1* gene is located on the plus strand, while the protein coding gene *VPS9D1* is located on the minus strand of human chromosome 16. NR045849 is a mouse lncRNA located on the plus strand near the *Vps9d1* gene (Figure S8A). When we ectopically expressed full-length NR045849 and VPS9D1-AS1 in MC38 and CT26W cell lines (Figure S8B), we observed an increase in the expression levels of TGF-β, TGFBR1, SMAD1 and STAT1 (Figure 6A). These findings indicated that NR045849 and VPS9D1-AS1 shared similar biological functions. Thus, we decided to explore the roles of VPS9D1-AS1 *in vivo* by OE in murine tumor cells.

**Fig. 6.**
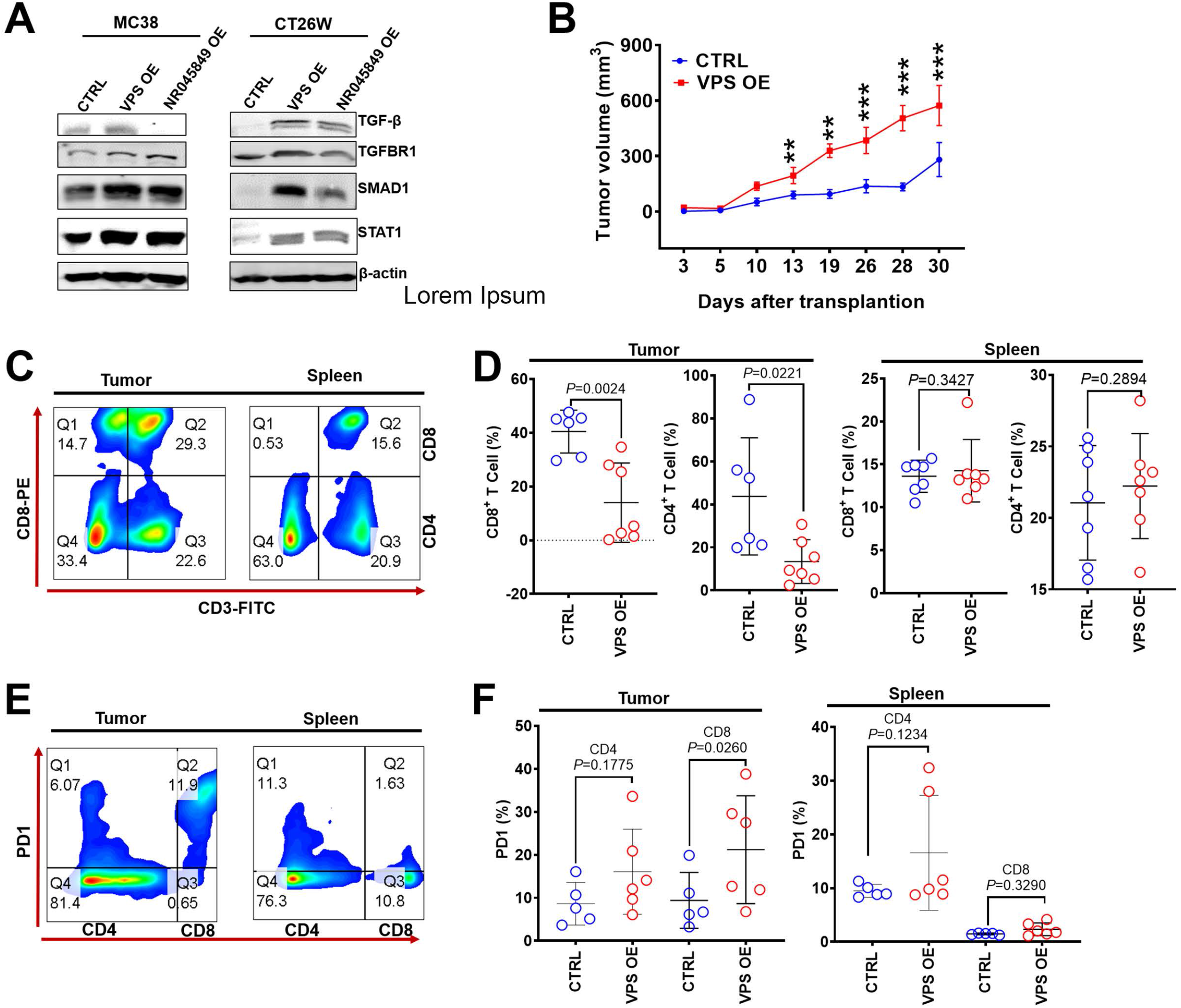
VPS9D1-AS1 OE cells inhibited T cell function *in vivo*. (A) VPS9D1-AS1 and NR045849 OE promoted the expression of TGF-β, TGFBR1, SMAD1, and STAT1. (B) Growth curves of MC38 xenograft tumors (n=7 per group). (C) Plots represent the percentages of CD8^+^ and CD4^+^ T cells in xenograft tumors and spleens. (D) Levels of CD8^+^ and CD4^+^ T cells were compared in the tumor and spleen. (E) FCM determined the PD1 levels in CD8^+^ and CD4^+^ T cells. (F) Levels of PD1 in CD4^+^ T and CD8^+^ T cells were compared between CTRL and VPS OE xenograft tumors (left) and spleens (right). *P* values were obtained by two-way ANOVA (B) and unpaired *t* nonparametric tests (D, F). Data are shown as the mean ± SEM (D, F). \*\**P*<0.01, \*\*\**P*<0.001. **Source data 1**. TGF-β, SMAD1, STAT1, TGFBR1and β-actin western-blot for Fig. 6A. **Figure supplement 8**. VPS9D1-AS1 inhibited TIL in xenograft tumors.

We further upregulated VPS9D1-AS1 more than 600-fold through three transfection cycles of lenti-VSP9D1-AS1 in MC38 cells, and VPS9D1-AS1 OE MC38 cells as well as control (CTRL) cells were subcutaneously injected into C57BL/6 mice (Figure S8C). MC38 VPS OE cell-derived tumors grew faster than MC38 CTRL cell-derived tumors (Figures 6B, S8D∼F). We also investigated T cells in mouse spleens and infiltrating T cells in tumor xenograft tissues (Figure 6C). The splenic CD8^+^ T and CD4^+^ T cells showed no significant differences between mice injected with MC38 VPS OE and CTRL cells (Figure 6D). When xenograft tumors were digested into single cells and analyzed by FCM, however, we found that VPS9D1-AS1 OE prevented CD4/8^+^ T cell from infiltrating into the tumor tissue (Figure 6D). These data underscore the inhibitory roles of VPS9D1-AS1 in T cell tumor infiltration.

We further assessed T cell surface markers that included CD44, CD107, CD62L, CCR7 and PD1 (Figures 6E and S8G). The levels of PD1 on CD8^+^ T cells was elevated in mice bearing MC38 VPS OE tumors (Figure 6F), while CD44, CD107, CD62L and CCR7 expression was not significantly different in either spleens or xenografted tumor tissues (Figure S8H). These results indicated that VPS9D1-AS1 regulated the pathway that controls the activation or differentiation of CD8^+^ T cells.

### VPS9D1-AS1 promotes tumorigenesis through Ifnar1 in AOM/DSS-induced intestinal cancer and acts as a therapeutic target

Using the VPS9D1-AS1 transgenic (VPS ^*Tg*^) mouse model, we generated an AOM/DSS model to further address the oncogenic roles of VPS9D1-AS1 *in vivo* (Figures S9A∼B). VPS ^*Tg*^ and wild-type (WT) mice from the same founder were subjected to AOM/DSS treatment cycles (Figure 7A). After AOM/DSS treatment, VPS ^*Tg*^ mice showed markedly higher intestinal tumor growth than WT mice (Figures 7B∼D and S9C). The VPS ^*Tg*^ mice had a shorter OS time than the WT mice (Figure 7E). Interestingly, VPS ^*Tg*^ mice showed lower CD8^+^ T cell infiltration (Figures 7F∼G). We further determined the levels of Ifnar1 in murine tumors and found that there were no differences in Ifnar1 levels between cancer and normal tissues (Figure S9D). However, Ifnar1 levels in tumor tissues were increased in VPS ^*Tg*^ mice (Figure 7H), indicating that the tumor-promoting effect of VPS9D1-AS1 OE was dependent on Ifnar1 expression.

**Fig. 7.**
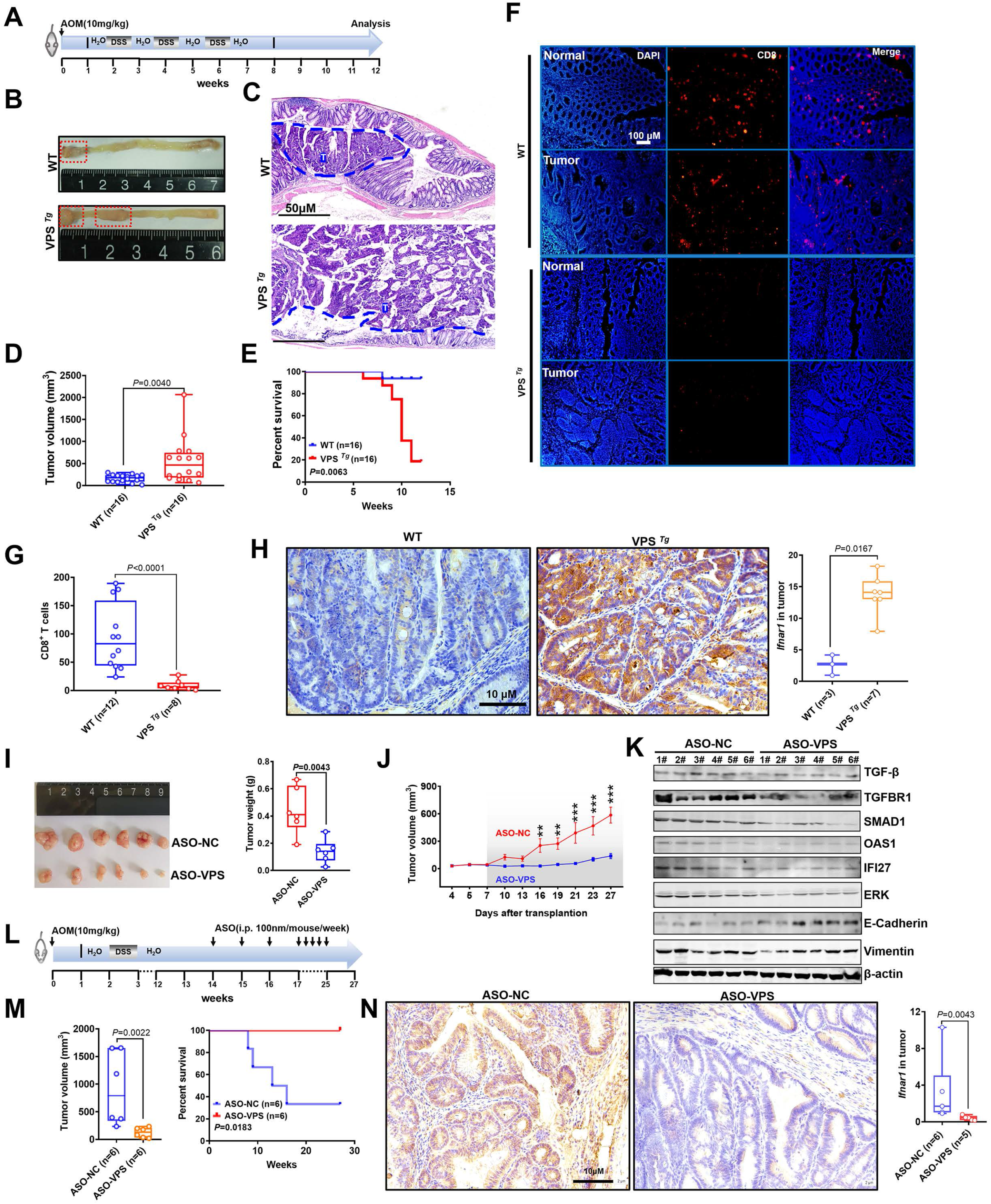
Transgenic mice validated the inhibitory role of VPS9D1-AS1 on CD8^+^ T cell infiltration and demonstrated the antitumor effects of targeting VPS9D1-AS1 with an antisense oligonucleotide (ASO) drug. (A) Treatment scheme of the AOM/DSS CRC model. Endpoint at 12^th^ weeks. (B) Representative images of colorectal tumors in mice of the indicated genotypes. (C) Hematoxylin (H)&eosin (E) staining showing the tumors (T). (D) Tumor volumes are plotted for WT and VPS ^*Tg*^ mice. (E) The OS curve depicts the difference in survival rate for WT and VPS ^*Tg*^ mice. (F) (G) VPS9D1-AS1 suppressed CD8^+^ T cell infiltration in AOM/DSS-induced CRC tissues. (H) The levels of Ifnar1 were upregulated in VPS ^*Tg*^ tumors compared with WT tumors. (I) Tumors, their weights and (J) growth curves of HCT116 xenograft tumors after injecting ASO-VPS (n=6) and ASO-NC (n=6). (K) Western blotting assays of the proteins involved in TGF-β, IFN, ERK, and EMT signaling in xenograft tumors. (L) Treatment scheme of ASO-treated mice with AOM/DSS-induced CRC. (M) ASO-VPS treatment decreased tumor volumes and increased OS of VPS ^*Tg*^ mice with AOM/DSS-induced CRC. (N) Ifnar1 expression upon ASO-VPS and ASO-NC treatment. *P* values were obtained by unpaired *t* nonparametric test (D, G, H, I, M, N), log-rank test (E, M), and two-way ANOVA tests (J). Data are shown as the mean ± SEM (D, G, H, I, M, N). \*\**P*<0.01, \*\*\**P*<0.001. **Source data 1**. SMAD1/5/9, PRS3, TGF-β and TGFBR1 down-western-blot for Fig. 7K. **Figure supplement 9**. VPS9D1-AS1 drives AOM/DSS-Induced mouse model of CRC.

To address the therapeutic potential of targeting VPS9D1-AS1, three 2-O-methyl antisense oligonucleotides (ASOs) specifically targeting VPS9D1-AS1 and one negative control were developed. Transfection of ASOs targeting VPS9D1-AS1 impaired the proliferation of HCT116, SW480 and MC38 VPS OE cells in culture (Figure S9E). Among them, ASO-siVPS3 showed the highest inhibition efficiency and was further used in an *in vivo* assay. We treated BALB/c nude mice bearing HCT116 xenograft tumors with intratumoral injection of ASO drugs and demonstrated that ASO-VPS significantly inhibited tumor growth *in vivo* (Figure 7I∼J and S9F). Moreover, the levels of TGF-β, TGFBR1, SMAD1, OAS1, IFI27, ERK, N-cadherin and vimentin were decreased, while E-cadherin was increased in ASO-VPS-treated xenograft tumors (Figure 7K).

VPS ^*Tg*^ mice were subjected to the AOM/DSS cycle to investigate whether VPS9D1-AS1 acts as a therapeutic target (Figures 7L and S9G). At the 12^th^ week, mice were divided into two groups and treated with ASO-NC and ASO-VPS drugs. In the ASO-NC-treated group, 4 (4/6) mice died because of aggressive tumor progression in the intestinal tract (Figure 7M). In contrast, all 6 mice treated with ASO-VPS survived until the final follow-up at the 27^th^ week. Among these 6 mice, 5 mice developed CRC, but their body weights were heavier than those of ASO-NC-treated mice from the 12^th^ week to the 27^th^ week (Figure S9H). ASO-VPS significantly reduced the tumor volumes (Figure 7M) and reduced Ifnar1 expression in murine tumors compared to ASO-NC treatment (Figure 7N). Collectively, our *in vivo* analyses indicated that VPS9D1-AS1 was the driver of CRC, inhibited CD8^+^ T cell infiltration and could serve as a therapeutic target.

## Discussion

Our present study shows that VPS9D1-AS1 synergizes with the TGF-β and IFN signaling pathways to form a signaling axis in which high intratumor cell activation prevents CTL evasion in the TME. Mechanistically, VPS9D1-AS1 enhances TGF-β signaling by increasing the expression of TGF-β, TGFBR1, and SMAD1/5/9 at the protein level through either ceRNA or scaffold mechanisms. Meanwhile, VPS9D1-AS1 KO in tumor cells downregulates the expression of many ISG genes, thus inactivating IFN signaling, and VPS9D1-AS1 KO in tumor cells increases the sensitivity of tumor cells to T cell cytotoxicity, which is illustrated by high levels of secreted IFN. Thus, VPS9D1-AS1/TGF-β/ISG signaling consists of the pathway that crosstalks between tumors and CD8^+^ T cells. Our *in vivo* assays support the use of VPS9D1-AS1 as a therapeutic target for CRC treatment (Figure 8).

**Fig. 8.**
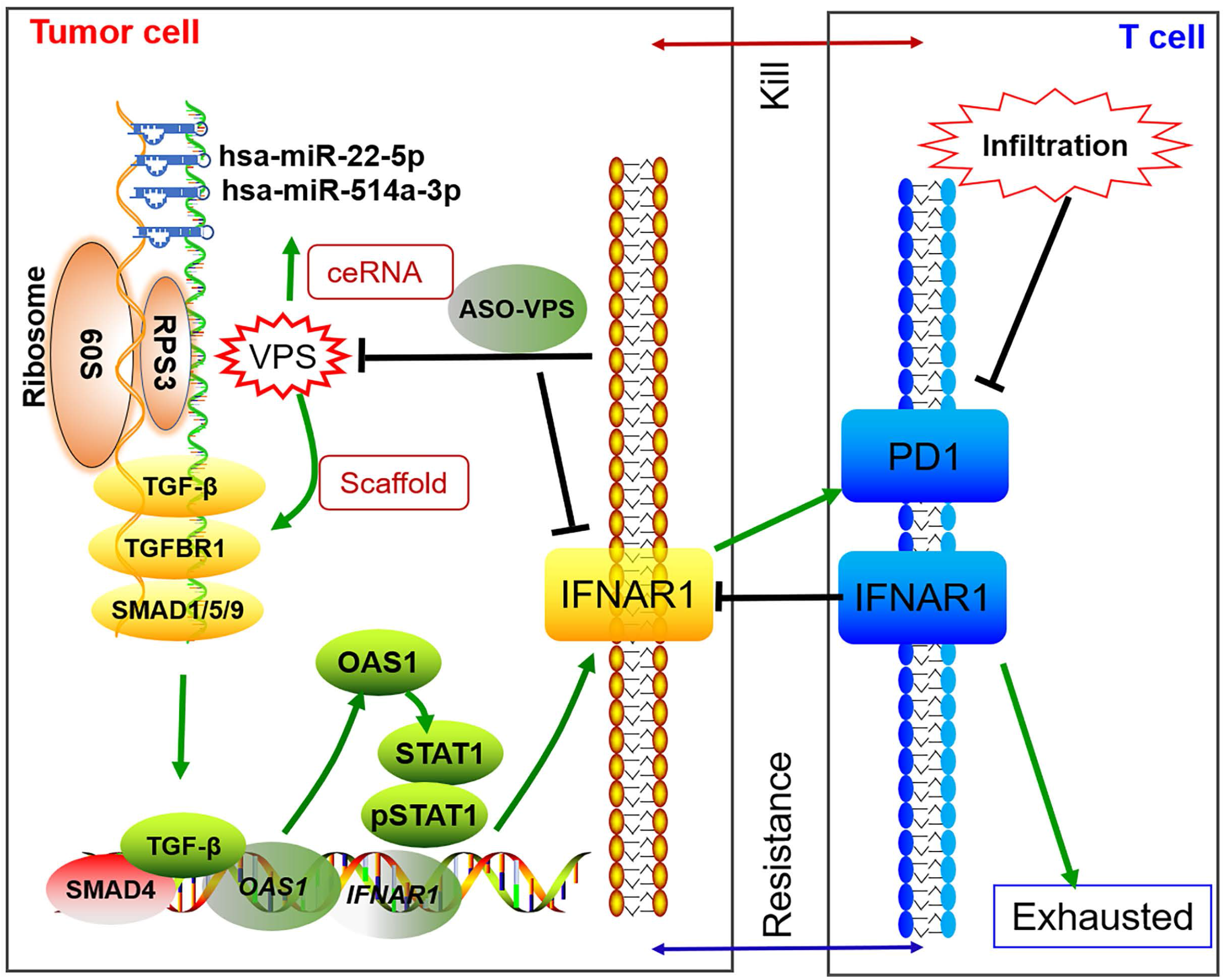
Illustration of the mechanism of VPS9D1-AS1 on promoting immune evasion via TGF-β signaling and ISGs (OAS1 and IFNAR1).

The dual roles of IFN signaling in tumor immune reactions have recently received intense attention (Benci et al., 2016). Autocrine type I IFNs from activated T cells augment T cell-mediated tumor regression (Chen et al., 2021) and are essential for the proliferation and differentiation of T cells (Lopes et al., 2021). In the TME, both tumor cells and infiltrating T cells express IFNA/GR, which sense and bind IFN molecules and then activate the expression of a series of ISGs (Grasso et al., 2021) (Jiang et al., 2018). On the other hand, the suppression of IFN signaling in tumor cells stimulates the production of chemokines, such as CXCL13, which results in tumor infiltration of NK cells and subsequent inhibition of tumor progression (Muthalagu et al., 2020). Our data show that VPS9D1-AS1 OE in CRC cells increases IFNAR1 through upregulated OAS1, which is an ISG gene. Hundreds of ISG genes have been identified, and they act either as tumor drivers, such as ADAR1 (Fritzell et al., 2019), or tumor repressors, such as UBA7 (Fan et al., 2020). We also found that both OAS1 and IFI27 are ISGs highly expressed in CRC. Consistent with these findings, OAS1 is a prognostic factor in breast cancer (Zhang and Yu, 2020), while IFI27 is overexpressed in cholangiocarcinoma (Chiang et al., 2019). In addition, OAS1 (but not IFI27) is the downstream ISG gene that is activated by VPS9D1-AS1.

IFN-activated STAT1 promotes PDL1 expression in tumors, which further accelerates tumor progression (Lv et al., 2021). This result is consistent with our finding that STAT1 is activated by VPS9D1-AS1. However, the effect of VPS9D1-AS1 on promoting IFN-induced phosphorylation of STAT1 does not persist for a long time and it might be associated with the role of pSTAT1 in activating the downstream cGAS-STING pathway, which is essential for defective-mismatch-repair-mediated immunotherapy (Guan et al., 2021).

Previous work has shown that TGF-β blocks IFN production, which is secreted in a paracrine fashion by activated CTLs and limits tumor regression (Guerin et al., 2019). Antibodies reduce TGF-β signaling in cancer stromal cells, facilitate T cell penetration into the center of tumors, and provoke antitumor immunity (Mariathasan et al., 2018). In the present study, we propose a model of autocrine signaling by cytokines in which VPS9D1-AS1 synergistically activates TGF-β and IFN signaling and induces cytokines in tumor cells to regulate CD8^+^ T cell infiltration and differentiate into an exhausted phenotype.

AOM/DSS-induced murine colorectal cancers frequently carry mutations in *Ctnnb1* and *Kras* and chronic inflammation-related MSI (Sharp et al., 2018), the Wnt pathway is constitutively activated, closely mirroring human CRC at the molecular level (Schulz-Heddergott et al., 2018). The *Tg* model we studied here highlights the driving role of VPS9D1-AS1, which might have been activated by mutations caused by AOM/DSS. Chronic inflammation contributes to intestinal symptoms, such as bleeding (Zhang et al., 2020). ASOs are chemically synthesized single-stranded oligonucleotides that selectively inhibit the target mRNA, and it is covalently linked to hepatocyte asialoglycoprotein receptor-binding N-acetylgalactosamine (GalNAc) to achieve cell-selective gene silencing (Yu et al., 2020). ASO drugs targeting VPS9D1-AS1 inhibited tumor-associated symptoms such as bleeding and prolonged survival time. Thus, this preclinical model demonstrates that suppressing VPS9D1-AS1 is an effective way to treat CRC.

In summary, we described the mechanism by which VPS9D1-AS1 promoted tumor immune evasion through the TGF-β signaling pathways and ISGs and provided compelling evidence that the VPS9D1-AS1/TGF-β/ISG axis might serve as a drug target to enhance the efficacy of ICB treatment against CRC. These findings expand our current mechanistic understanding of CRC progression and provide potential therapeutic approaches by targeting VPS9D1-AS1 to enhance immunotherapy in patients with CRC.

## Materials and methods

Detailed procedures are provided in the Supplemental Materials and Methods.

### RNAScope *In Situ* Hybridization Assay

RNA *in situ* hybridization assays were performed using an RNAscope kit (Advanced Cell Diagnostics, Hayward, CA, USA). VPS9D1-AS1 mRNA molecules were detected with single-copy detection sensitivity. Single-molecule signals were quantified on a cell-by-cell basis by manual counting. The signals per cell were evaluated to quantitate the levels of VPS9D1-AS1. The signals were graded as 0 (0-1 dots/10 cells), + (1-3 dots/cell), ++ (4-10 dots/cell), +++ (> 10 dots/cell with <10% of dots in clusters), and ++++ (> 10 dots/cell with >10% of dots in clusters) (Zhao et al., 2018). The scores were classified as follows: “-, +” represented negative expression and “++, +++, ++++” represented positive expression of VPS9D1-AS1.

### Mice

Conditional floxed human VPS9D1-AS1 knock-in alleles with LoxP sites were introduced into the *Rosa26* gene locus to construct C57BL/6J-Gt (ROSA)26Sor^em(CAG-VPS9D1-AS1)1Smoc^ mice. A mixture of the construct and CRISPR/Cas9 vector was microinjected into zygotes. The zygotes were implanted into foster mice. The above procedures were performed by Shanghai Model Organisms. Successful integration in the founder mice was identified by PCR analyses of genomic DNA using primers targeting VPS9D1-AS1. After screening, the positive founder (bearing R26-eCAG-VPS9D1-AS1) was crossed with Cre-*villin* mice to produce conditional VPS9D1-AS1 transgenic (VPS ^*Tg*^) mice in the intestinal epithelium.

To induce colorectal cancer with AOM/DSS, C57BL/6 VPS ^*Tg*^ mice received a single intraperitoneal injection of azoxymethane (AOM) (Sigma Aldrich) at a dose of 10 mg/kg body weight. One week later, animals were exposed to 1∼3 cycles of 2% dextran sulfate sodium salt (DSS, MW=36,000-50,000, MP Biomedicals, CAT# 216011050). For *in vivo* inhibition of VPS9D1-AS1, mice in the treatment group received 10 nmol (100 μl, 100 mM ASO in PBS) ASOs *i*.*p*., once a week. Intestinal tumor volumes were calculated by the formula: π × (1/2 width)^2^ × length.

For the xenograft tumor model, 2 × 10^6^ HCT116 or SW480 cells were implanted subcutaneously in 6-week-old female BALB/c nude mice (Charles River). For MC38 cells, 4 × 10^6^ cells were subcutaneously injected into the lower abdominal region of 6-week-old C57BL/6 wild-type mice (Charles River). Tumors were assessed 2∼3 times a week by a caliper measurement, and the volumes were calculated with the following formula: 1/2 × (length × width^2^). MC38 cells were stably transfected with luciferase to label tumor progression and were monitored using a Carestream *in vivo* imaging system (MS FX Pro). Animal experimental protocols were approved (AEEI-2021-105) according to the guidelines of the Ethics Committee for Animal Testing of Capital Medical University.

### Multispectral fluorescence immunohistochemistry (mfIHC)

For quantitative evaluation of the infiltrating lymphocytes, multispectral imaging was employed to stain samples with antibodies on a TMA slide. Multispectral fluorescence immunohistochemistry (mfIHC) assays were performed using a PerkinElmer Opal™ 7-color mfIHC Kit (PerkinElmer) according to the manufacturer’s instructions.

### Statistical analysis

Statistical analysis was performed using GraphPad Prism software and R software. Student’s *t* test or analysis of variance (ANOVA) (one- or two-way) with *Bonferroni* post hoc test were used to evaluate the statistical significance. The survival curves were plotted according to the Kaplan-Meier method and evaluated by the log-rank test. A *P* value less than 0.05 was considered statistically significant.

## Abbreviations

VPS9D1-AS1: VPS9 domain-containing protein 1 antisense RNA 1;
Cre: cyclization recombination;
EMT: Epithelial mesenchymal transition;
STAT1: Signal transducer and activator of transcription;
cGAS: Cyclic GMP-AMP Synthase;
STING: Stimulator of interferon genes protein;
AOM: Azoxymethane;
DSS: Dextran sodium sulfate;
SMAD: Mothers against decapentaplegic homolog;
PD1: Programmed death-1;
PDL1: Programmed death ligand 1.

## Acknowledgments

We thank Dr. Jian Liu for providing CRISPR/Cas9 virus packaging vectors and Dr. Suliang Guo for his assistance in animal feeding.

## Author’ Contributions

LY designed the experiments and wrote the manuscript. LY, HQ, XXH and JJT performed the experiments. HQ and ZJW collected patient samples. LY and TW acquired funding. All authors read and approved the final manuscript.

## Ethics approval

This study was approved by the Institutional Review Board at Beijing Chao-yang Hospital. All enrolled subjects signed written informed consent forms. All animal studies were conducted with the approval of the Institutional Animal Care and Use Committee of Capital Medical University.

## Data and materials availability

The data that support the findings of this research are available, deposited in Sequence Read Archive (accession number: PRJNA716724) and Dryad (URL: https://datadryad.org/stash/share/i7c2Q5C9qLIvEFZD8_F0jKD8bOJ_dXaWPqRRQ3FCiu4).

## Supplemental Figure Legends

**Figure S1.**
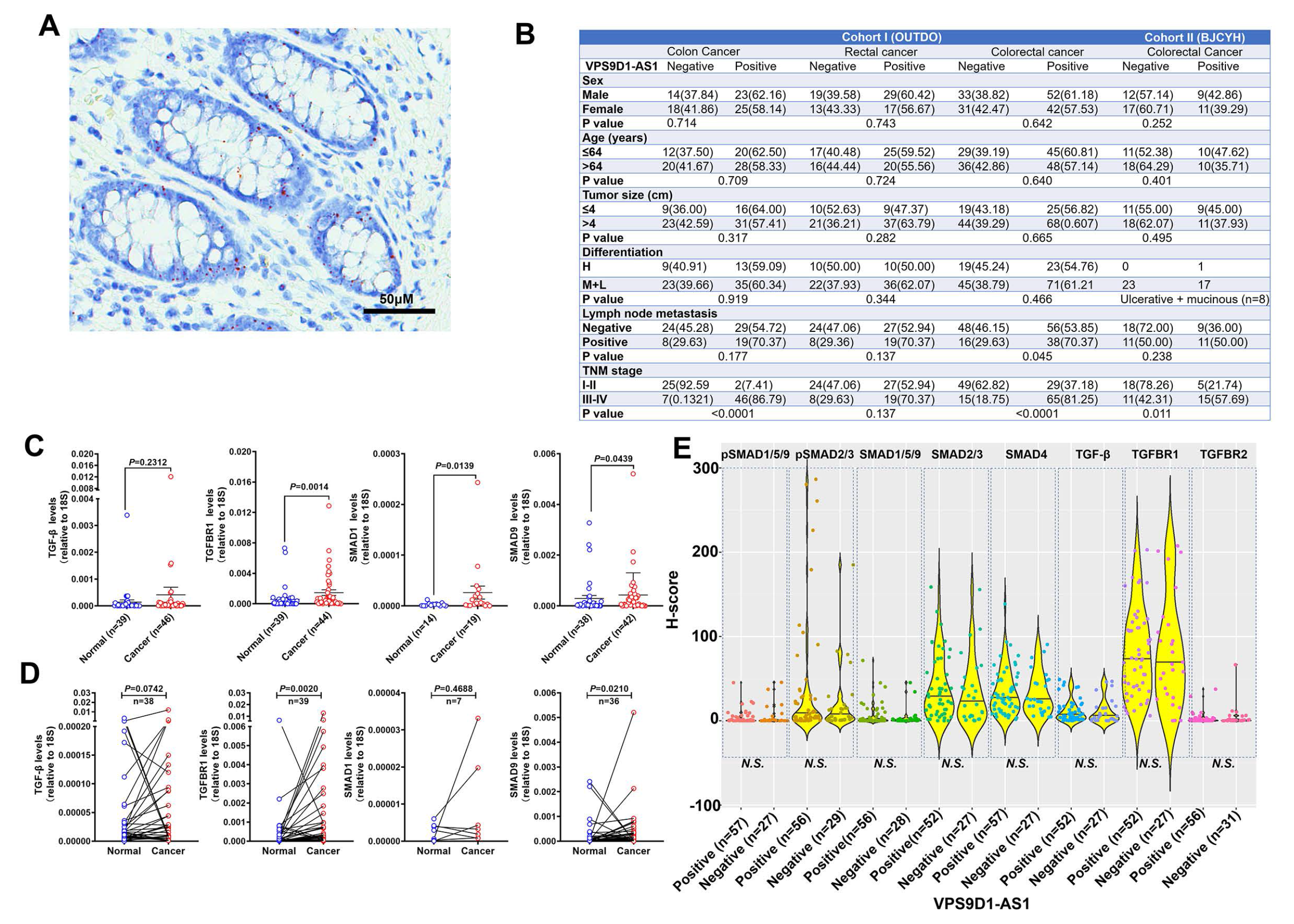
Levels of VPS9D1-AS1 were not related to TGF-β signaling in cancer stromal cells. (A) Representative image of VPS9D1-AS1 expressed in normal colonic epithelial cells. (B) Clinical pathologic analyses demonstrated that VPS9D1-AS1 was correlated with lymph node metastasis and TNM stage in the OUTDO and BJCYH cohorts. (C, D) qRT-PCR was used to determine the mRNA levels of TGF-β, TGFBR1, SMAD1, and SMAD9 in the BJCYH cohort. (E) There were no significant relationships between VPS9D1-AS1 levels and TGF-β signaling in cancer stromal cells in OUTDO cohort. *P* values were obtained by *Wilcoxon* rank-sum test (B), unpaired *t* nonparametric test (C) and paired *t* nonparametric test (D). Data points are presented as the mean ± SEM (C) and the minimum, first quartile, median, third quartile, and maximum (E). *N*.*S*. not significant.

**Figure S2.**
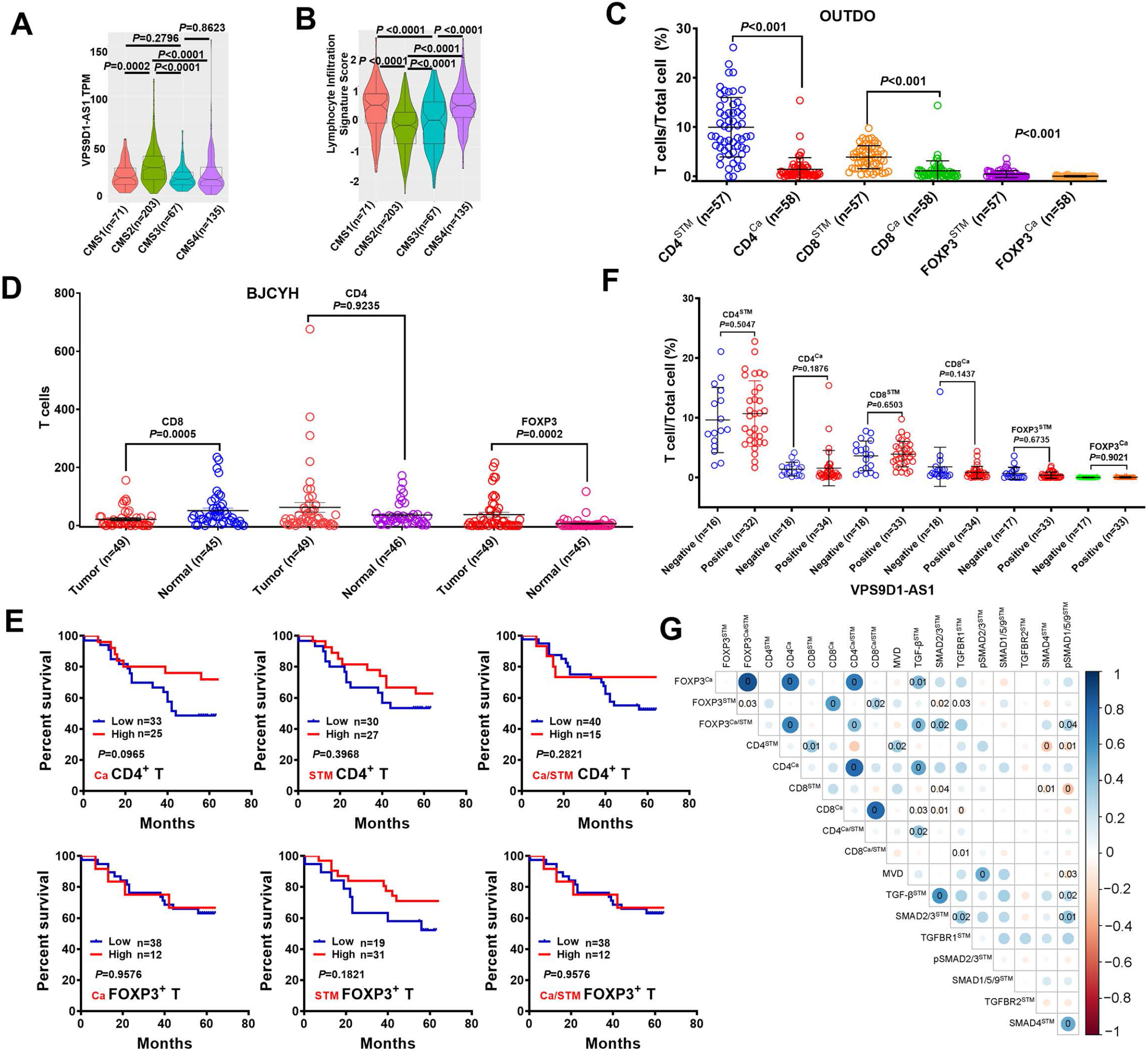
Integrative analysis of the relationship between VPS9D1-AS1, TGF-β signaling and TILs. (A) TCGA analysis confirmed that the highest VPS9D1-AS1 expression was the predominantly in CMS2-type CRC patients. (B) CMS2 patients showed the lowest lymphocyte infiltration signal score. (C) Comparison of the percentages of CD4^+^, CD8^+^, and FOXP3^+^ T cells in cancer and cancer stromal tissues of OUTDO cohort. (D) Comparison of the number of CD4^+^, CD8^+^, and FOXP3^+^ T cells in cancer and normal tissues of the BJCYH cohort by IHC assays. (E) Kaplan-Meier OS curves showed that CD4^+^ T and FOXP3^+^ T cells had no prognostic significance. (F) Proportions of CD4^+^, CD8^+^, and FOXP3^+^ T cell out of the total cells of Ca and STM tissues were compared according to the levels of VPS9D1-AS1. (G) *Pearson* correlation analyses investigated the relationships between TILs and TGF-β signaling in cancer stromal tissues. *P* values were obtained by unpaired *t* nonparametric test (A, B, C, D, F), log-rank test (E), and *Pearson* correlation test (G). Data are shown as data points with mean ± SEM (C, D, F).

**Figure S3.**
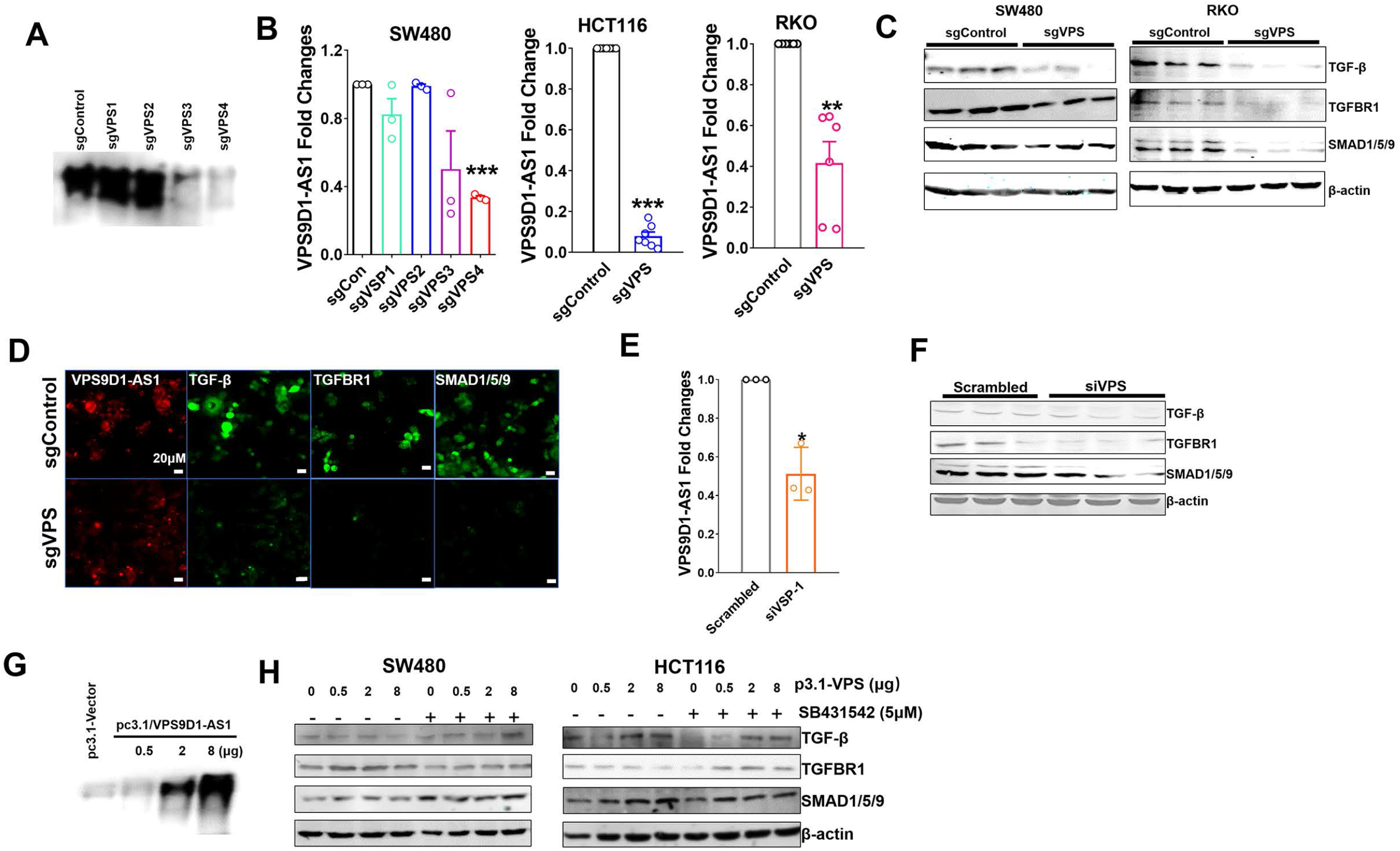
VPS9D1-AS1 activated TGF-β signaling. (A) Northern blotting detected the effectiveness of sgRNA targeting VPS9D1-AS1 in SW480 cells. (B) qRT-PCR was used to determine the levels of VPS9D1-AS1 in stable KO cells (sgControl vs. sgVPS). (C) Western blotting identified the levels of TGF-β, TGFBR1, and SMAD1/5/9. (D) RNA FISH-IF showed the levels of VPS9D1-AS1 and proteins that included TGF-β, TGFBR1, and SMAD1/5/9 in HCT116 cells. (E) qRT-PCR showed the downregulation of VPS9D1-AS1 after siRNA (siVPS) transfection in HCT116 cells. (F) Changes in TGF-β, TGFBR1, and SMAD1/5/9 after siRNA transfection in HCT116 cells. (G) Northern blotting confirmed the overexpression of VPS9D1-AS1 by transfection with the pcDNA3.1 vector in SW480 cells. (H) Effects of VPS9D1-AS1 overexpression (OE) on TGF-β, TGFBR1, and SMAD1/5/9 in SW480 cells. *P* values were obtained by paired *t* tests (B, E). Data are shown as the mean ± SEM (B, E). \**P*<0.05, ** *P*<0.01, *** *P*<0.001.

**Figure S4.**
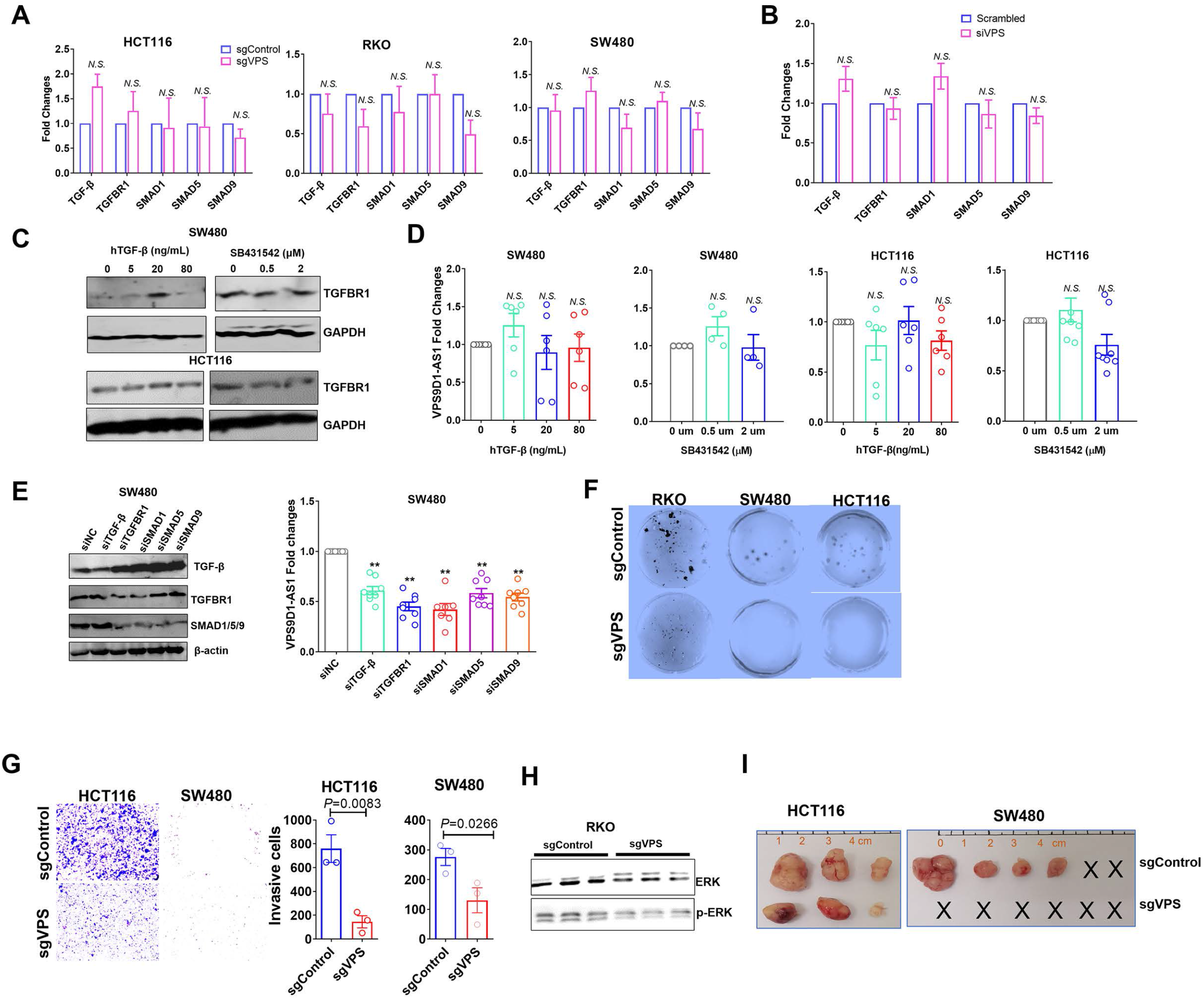
VPS9D1-AS1 regulated TGF-β signaling and promoted tumor proliferation and migration. (A, B) mRNA levels of *TGF-β, TGFBR1, SMAD1, ∼5, and ∼9* after VPS9D1-AS1 KO or KD. (C) Human recombinant (h) TGF-β- and SB431542-treated SW480 and HCT116 cells. (D) hTGF-β and SB431542 had no effect on VPS9D1-AS1 levels. (E) VPS9D1-AS1 levels were decreased by siTGF-β, siTGFBR1, and siSMAD1, ∼5, ∼9. (F) Clone forming assay results of RKO/SW480/HCT116 sgControl and sgVPS cells after culturing for 14 days. (G) Transwell assays determined the migration of HCT116 cells. (H) The levels of ERK and pERK in RKO cells (the interference gene was same with used as in Figure S3C). (I) HCT116/SW480 sgControl and sgVPS cells were separately transplanted subcutaneously into BALB/c nude mice. *P* values were obtained by paired or unpaired *t* tests (A, B, D, E, G). Data are shown as the mean ± SEM (A, B, D, E, G). *N*.*S*. not significant, ** *P*<0.01.

**Figure S5.**
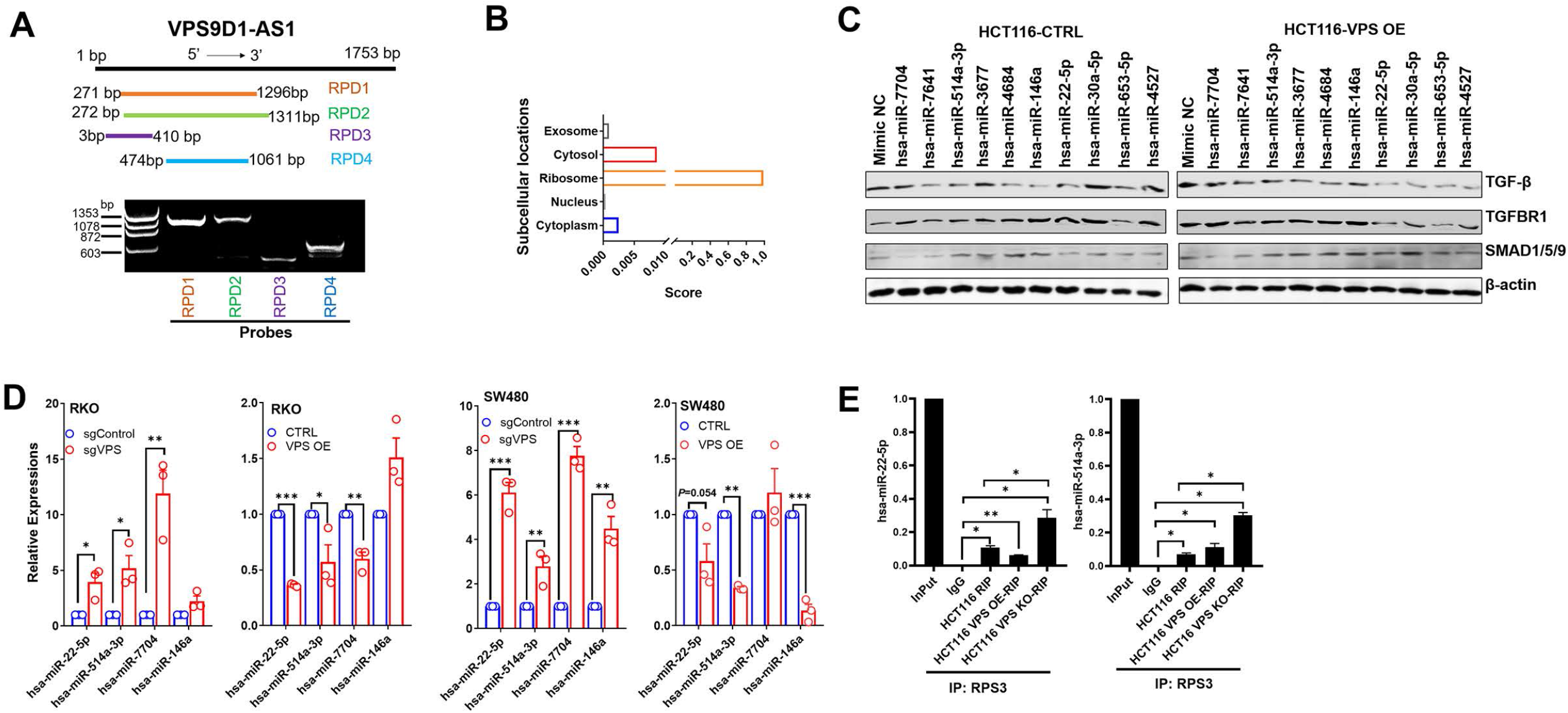
VPS9D1-AS1 functions as the scaffolding lncRNA and ceRNA. (A) Synthesized probes used in RNA pulldown (RPD) assays for targeting the sequence of VPS9D1-AS1. (B) Subcellular localizations of VPS9D1-AS1 were analyzed by online tools (www.csbio.sjtu.edu.cn/bioinf/lncLocator/). (C) Western blotting was used to analyze the levels of TGF-β, TGFBR1, and SMAD1/5/9 after transfecting 10 miRNA mimics into HCT116 CTRL and VPS OE cells. (D) qRT-PCR validated the differential expression of hsa-miR-22-5p, hsa-miR-514a-3p, hsa-miR-7704, and hsa-miR-146a in RKO and SW480 cells. (E) RIP assay determined the interaction between RPS3 and hsa-miR-22-5p and hsa-miR-514a-3p. *P* values were obtained by paired *t* tests (D, E). Data are shown as the mean ± SEM (D, E). \**P*<0.05, ** *P*<0.01, *** *P*<0.001.

**Figure S6.**
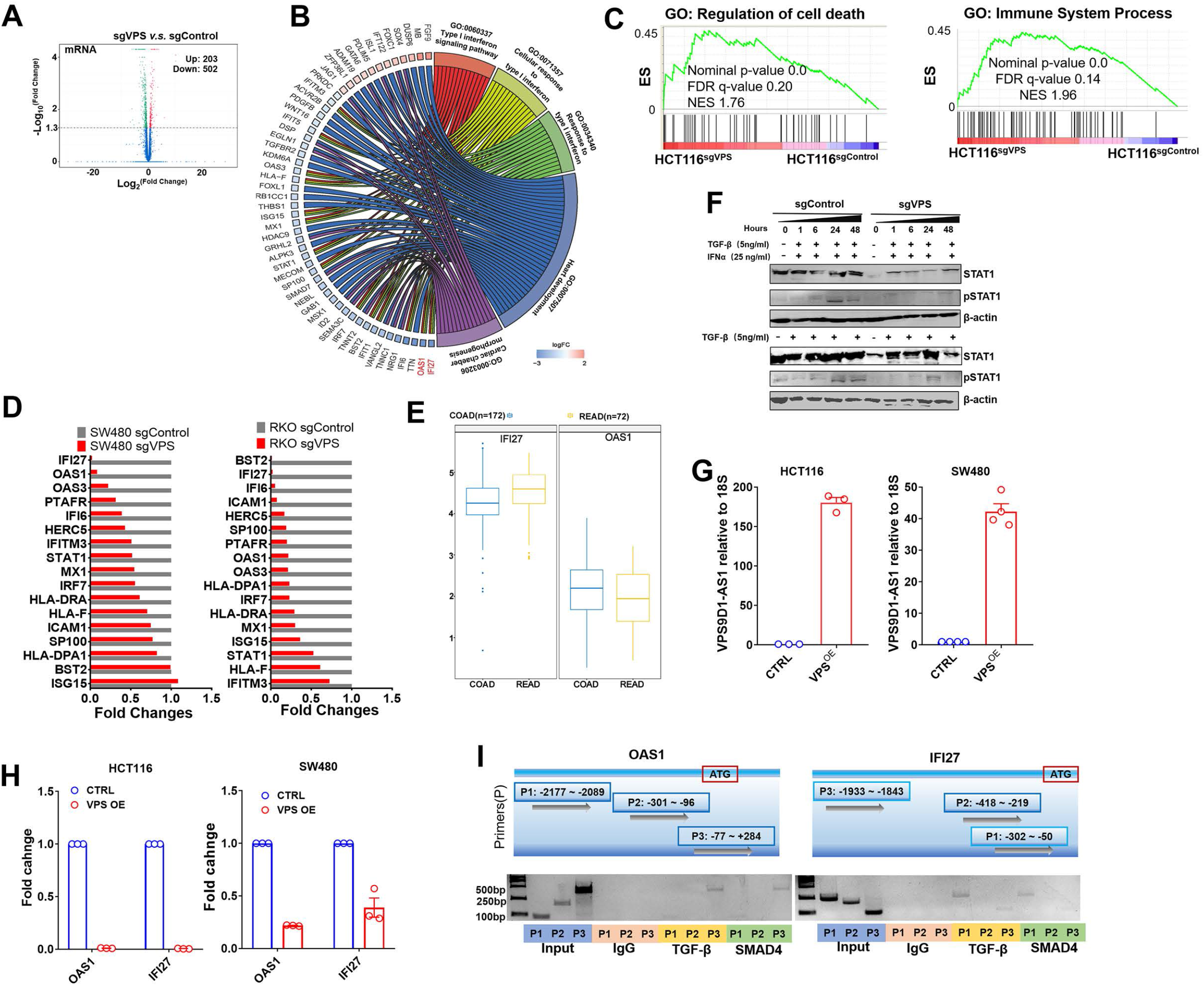
VPS9D1-AS1 plays a role on interferon signaling. (A) Volcano plot displaying the differential mRNA profiles of HCT116^sgControl^ and HCT116^sgVPS^ cells. (B) Gene Ontology analyses explored the roles of VPS9D1-AS1 in regulating IFNα/β signaling. (C) GSEA revealed that VPS9D1-AS1 was involved in pathways related to cell death and immune system processes. (D) Differential expression of 17 proteins in the IFNα/β signaling pathway was validated in RKO and SW480 cells. (E) TCGA COAD and READ datasets confirmed the overexpression of OAS1 and IFI27. The fold changes for OAS1 and IFI27 expression in cancer tissues relative to normal tissues were shown. (F) hTGF-β prevents hIFNα from activating STAT1 phosphorylation. (G) Fold changes of VPS9D1-AS1 in HCT116 and SW480 VPS9D1-AS1 OE stale cell lines relative to CTRL stale cell lines. (H) Levels of IFI27 and OAS1 in cells with VPS9D1-AS1 overexpression. (I) ChIP assays determined the interaction of antibodies against TGF-β and SMAD4 with the promoter regions of *OAS1* and *IFI27* in HCT116 cells. Data are shown as the mean ± SEM (G, H).

**Figure S7.**
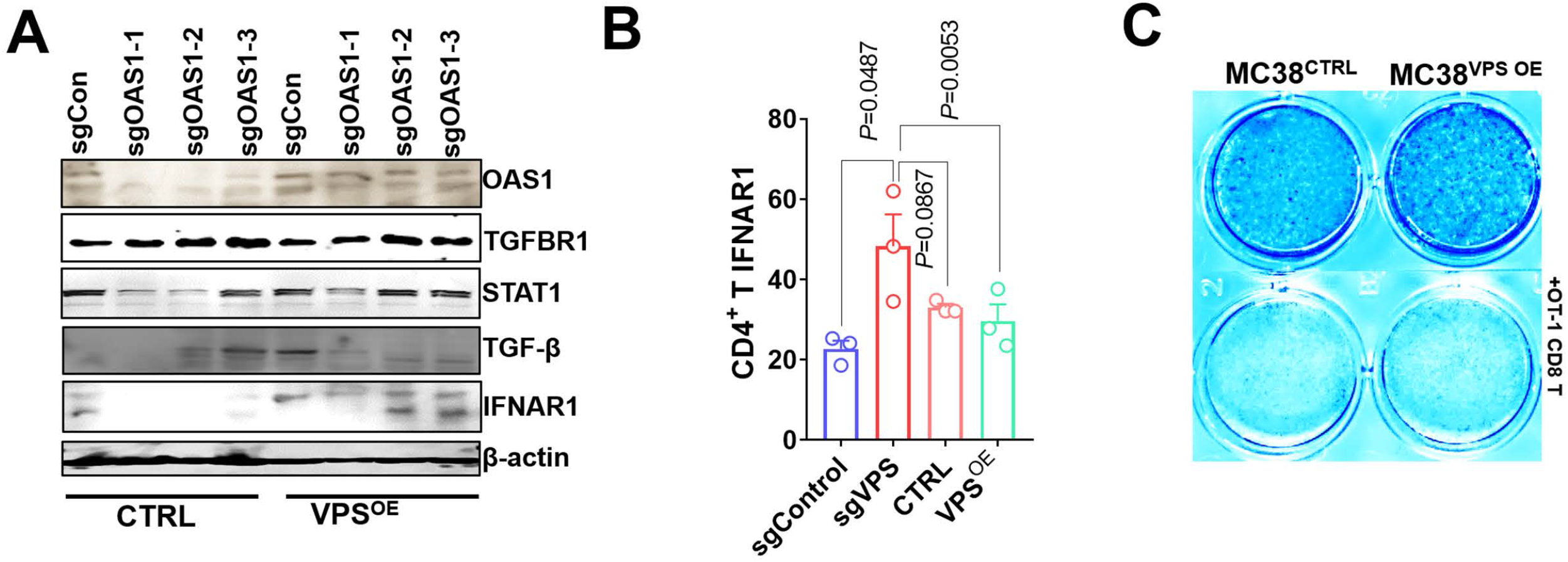
IFNAR1 mediates crosstalk between T cells and cancer cells. (A) Western blotting assay to determine the levels of OAS1, TGFBR1, STAT1, TGF-β, and IFNAR1 in HCT116 cells. (B) IFNAR1 levels in CD4^+^ cells after coculturing with sgControl, sgVPS, CTRL, and VPS OE cells. (C) Representative results of T cell cytotoxicity assays of the indicated MC38^CTRL^ and MC38^VPS OE^ cell lines after exposure to OT-1 CD8^+^ T cells. *P* values were obtained by unpaired *t* test (B). Data are shown as the mean ± SEM (B).

**Figure S8.**
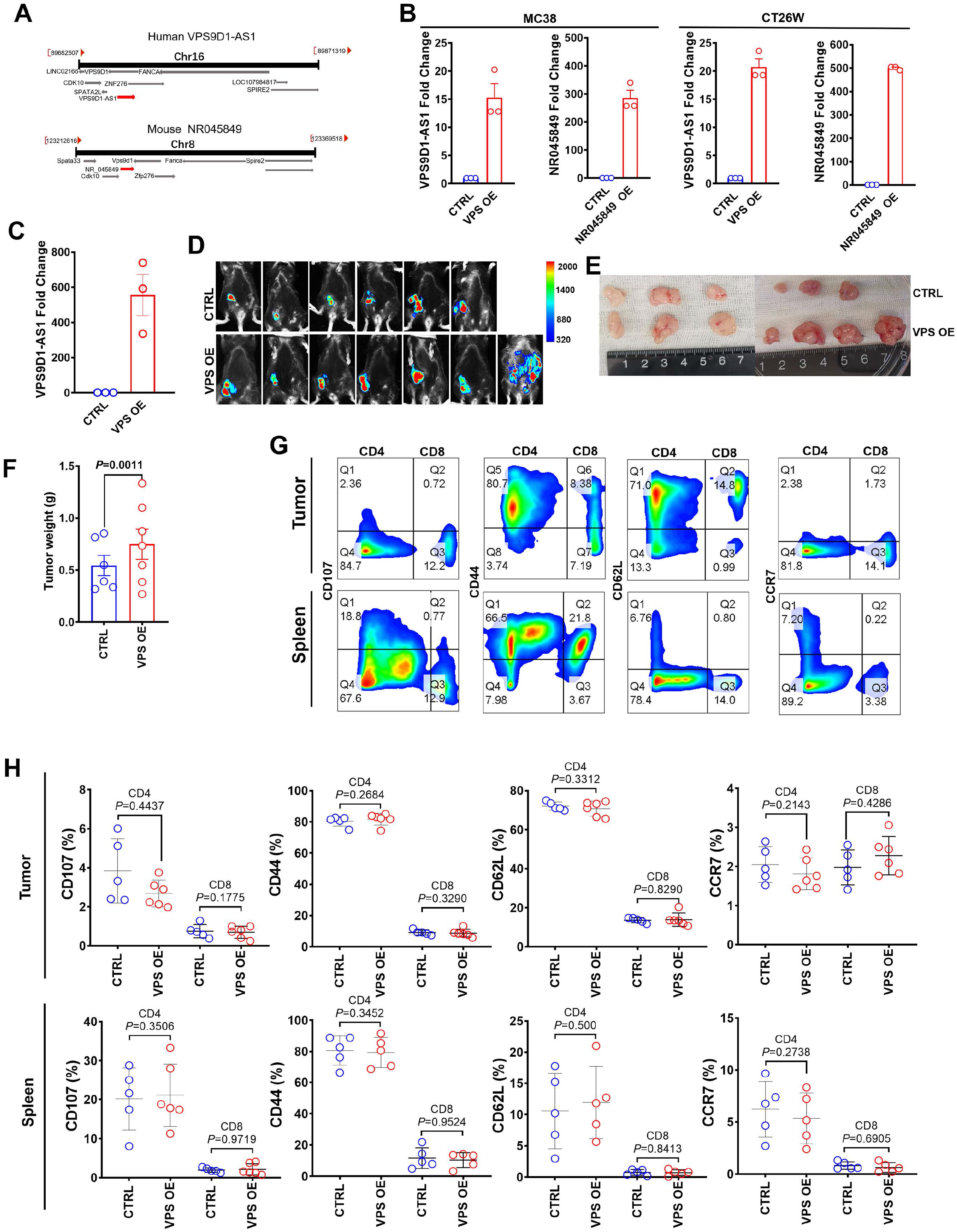
VPS9D1-AS1 inhibited TIL in xenograft tumors. (A) Schematic of human VPS9D1-AS1 and murine NR045849 locations on the chromosome. (B) Fold changes in the expression levels of VPS9D1-AS1 and NR045849 in MC38 and CT26W cells. (C) qRT-PCR confirmed the overexpression of VPS9D1-AS1 after three transfections with lentivirus vectors in MC38 cells. (D) *In vivo* imaging showed transplanted MC38^CTRL^ and MC38^VPS-OE^ xenograft tumors. (E) Xenograft tumors were harvested 35 days after injection (one mouse in the CTRL group did not form tumors). (F) Xenograft tumor weights for MC38^CTRL^ and MC38^VPS OE^. (G) Representative FCM results of CD44, CD62L, CD107, and CCR7 in CD4^+^ and CD8^+^ T cells. (H) Comparisons of the levels of CD44, CD62L, CD107, and CCR7. *P* values were obtained by unpaired *t* test (F, H). Data are shown as the mean ± SEM (F, H).

**Figure S9.**
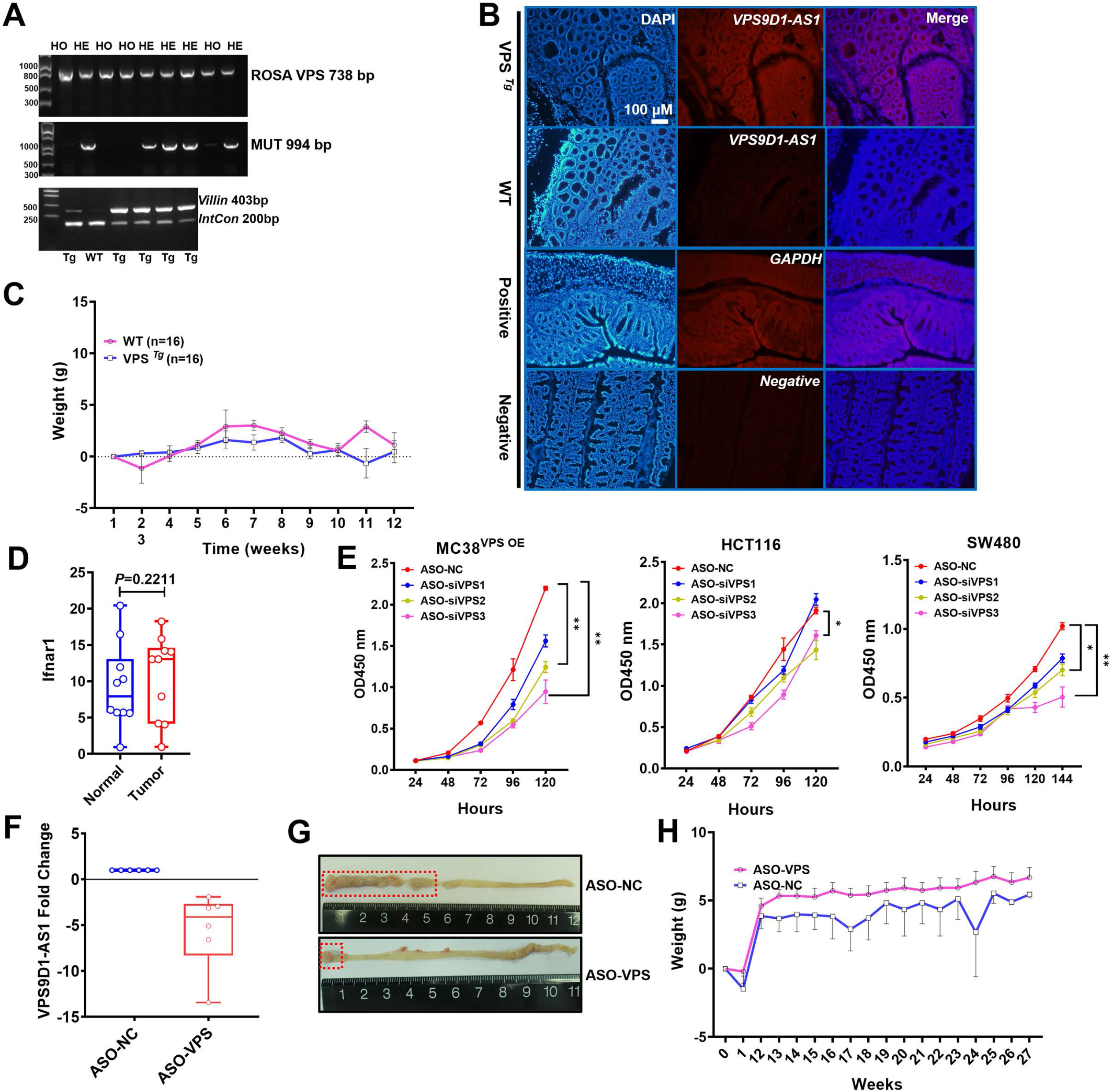
VPS9D1-AS1 drives AOM/DSS-Induced mouse model of CRC. (A) Genotyping of VPS ^*Tg*^ mice. Internal Control (IntCon). (B) RNA FISH identified VPS9D1-AS1 expression in VPS ^*Tg*^ and wild-type (WT) mice. A probe targeting *GAPDH* mRNA was used as a positive control. (C) Body weight changes upon AOM/DSS treatment. (D) Comparison of Ifnar1 expression in normal tissue and tumor tissue. (E) CCK-8 assays for evaluating the proliferation of MC38^VPS OE^, HCT116, and SW480 cells after ASO treatment. (F) Fold changes in the levels of VPS9D1-AS1 in xenograft tumors. (G) Representative pictures of the mouse intestine treated with ASO-NC and ASO-VPS. (H) VPS ^*Tg*^ mice were treated with ASO-VPS and ASO-NC, and body weight changes were measured weekly. *P* values were obtained by one-way ANOVA (E) or unpaired *t* nonparametric test (D). Data are shown as the mean ± SEM (C, E, H) or box plots with minimum, first quartile, mean, third quartile, and maximum values (D, F).

